# SLUR(M)-py: A SLURM Powered Pythonic Pipeline for Parallel Processing of 3D (Epi)genomic Profiles

**DOI:** 10.1101/2024.05.18.594827

**Authors:** Cullen Roth, Vrinda Venu, Sasha Bacot, Shawn R. Starkenburg, Christina R. Steadman

## Abstract

Epigenomics has become multi-faceted, with researchers exploring chromatin structure, nucleosome states, and epigenetic modifications, producing large, complex multi-omic data sets. Given this shift, there is demand for bioinformatics that leverage high performance computing (HPC) and parallelization to quickly process data. As such, we developed SLUR(M)-py: a pythonic computational platform that leverages the Simple Linux Utility for Resource Management system (SLURM) to process sequencing data. SLUR(M)-py is multi-omic and automates calls to SLURM for processing paired-end sequences from chromatin characterization experiments, including whole-genome, ChIP-seq, ATAC-seq, and Hi-C, thereby eliminating the need for multiple analytics pipelines. To demonstrate SLUR(M)-py’s utility, we employ ATAC-seq and Hi-C data from viral infection experiments and the ENCODE project, and illustrate its processing speed and completeness, which outpaces current HPC pipelines. We explore the effect of dropping duplicate sequenced reads in ATAC-seq, demonstrate how SLUR(M)-py can be used for quality control, and how to detect artifacts in Hi-C from viral infection experiments. Finally, we show how features in SLUR(M)-py, like inter-chromosomal analysis, can be used to explore the dynamics of chromosomal contacts in mammalian cells. This multi-omic, system agnostic platform eases the computational burden for researchers and quickly produces accurate, reliable data analytics for the epigenomics community.

## Introduction

Within the nucleus, the shape, looping, modification, and interactions between chromatin are critical to cellular function, development, and for mounting responses to environmental stimuli [1–14]. A myriad of sequencing technologies, based on paired-end, short read sequencing, are designed to probe various characteristics of chromatin, including conformation, chemical modification, and resulting gene expression. Assays including chromatin conformation capture followed by sequencing (Hi-C) and for transposase-accessible chromatin with sequencing (ATAC-seq) are used to study the three-dimensional (3D) architecture and the accessibility of DNA within cells, respectively [15–19]. Additionally, chromatin immunoprecipitation (ChIP-seq) assays probe the genome for specific chemical modifications made to histone proteins, simplistically indicative of permissive or repressive chromatin states [20–22]. Together, each of these sequencing paradigms provide complementary information important to researchers when exploring a hypothesis pertaining to epigenetics. Thus, the ability to quickly integrate, characterize and standardize data from these sequencing approaches in epigenomic investigations (including their complete processing, analysis, and visualization) is critical to understand chromatin dynamics during cellular development and responses to external stimuli.

While each genomic sequencing strategy has specific benefits, caveats, and requirements for analysis, the expected output of most genome-wide sequencing assays is the same: paired-end sequencing data. From the perspective of a bioinformatician, across experiments and assays, the processing of paired-end sequencing data is similar between the aforementioned genomic and epigenomic assays. Processing of both types of data requires trimming of adaptors from reads, removal of low-quality reads (such as reads with repetitive, low-complexity sequence), alignment to a reference genome, marking and removal of duplicate alignments, filtering of poorly aligned or non-unique alignments, and removal of artifacts. As such, co-processing these data through a single informatics pipeline can simplify downstream analysis, visualization, and biological interpretation. Individual single-omic software tools and pipelines have been written for this purpose. For example, Hi-C data is traditionally analyzed through the Juicer pipeline, which has automated the analysis and generation of interacting chromatin maps (.hic files) from paired-end sequencing files to characterize chromatin loops and topological associated domains (TADs) [23, 24], both important features to report when presenting Hi-C data. HiCExplorer and Fan-c, which are python based and have command line functionality, offer the research community alternative pipelines for generating Hi-C maps, data analysis, and visualization [25–27]. For epigenomic assays like ATAC-seq and ChIP-seq, the ENCODE project has guidelines and recommended software for experiments and subsequent generation of sequencing files (.bam) for appropriate, high-quality analysis [18,22]. Additionally, software such as ATAC-graph provides automated processing, diagnostics, and quality control plots of ATAC-seq data [28]. Investigators will often cobble these tools together with other bioinformatic tools to analyze the myriad of sequencing data that is generated within a single study [29, 30]; while sound, this can be challenging, time consuming, inefficient, and fraught with the potential for (human) error. Moreover, the amalgamation of disparate pipelines for analysis of data from the same set of experiments can generate inconsistencies and incompatibilities.

One feature of many of these pipelines is their “Pythonic” nature, that is, many are written in Python. Python is an ideal coding language to process paired-end sequencing reads from epigenomic and chromatin conformation experiments given its low barrier to entry, flexibility, and open-source nature. While these pipelines leverage Python—and are therefore accessible—they are only equipped to process single omic data types. Further, in many instances, these pipelines do not scale with larger data sets. As such, using these tools to process numerous ‘omics data sets that can be quite large (GB) from human cell lines or model systems can prove problematic, when more than one billion reads are recommended for fully resolved genomes [4, 31].

To analyze large cohorts of sequencing data, like Hi-C from several experiments, or multi-layered omics data, researchers regularly rely on a high-performance computing (HPC) environment to quickly process data; these include popular HPC environments such as Amazon Cloud Computing and Nextflow [32]. Alternatively, web-based applications have also been developed for the processing of sequencing data [25,33–36]. One of the most popular open-source software tools, designed to manage a high-performance computing environment is the Simple Linux Utility for Resource Management system (SLURM) [37]. As its name implies, SLURM is simple to use and scalable for large computational jobs [38]. Given these features and the user-friendly nature of SLURM, it has become an accepted and integral tool for HPC, capable of managing small to large servers. SLURM empowers users to submit and run parallel jobs while also establishing dependencies, allowing for the creation of truly end-to-end, automated pipelines [38]. For example, the Juicer pipeline leverages SLURM to manage, submit, and run parallel processes for generating .hic files from Hi-C experiments [23]. Additionally, the Hi-C-Pro pipeline contains a parallel mode which can submit jobs to SLURM to create and run parallel bioinformatic tasks [39]. To our knowledge, there is currently no other published HPC enabled pipeline for the processing and analysis of most other sequencing data from chromatin characterization experiments such as whole-genome, ChIP-seq, or ATAC-seq.

To fill this gap, we combine the accessibility of Python with the utility and scalability of the SLURM environment to provide the epigenomics community with a flexible pipeline for analyzing large (GB) multi-omics data sets. We call it SLUR(M)-py (pronounced slurpy; the M is silent): this Python-based pipeline is designed for a HPC environment that processes the alignments, filtering, and analysis of various types of paired-end sequencing data (genomics and epigenomics). This Pythonic pipeline auto-generates and submits scripts to SLURM; SLURM then schedules and parallelizes bioinformatic processes underlying the analysis of ‘omics data, greatly improving the speed of common bioinformatic tasks (by a factor of eight). The novelty of SLUR(M)-py is its ability to process paired-end sequencing data from several *different* sequencing strategies, including whole-genome, ATAC-seq, ChIP-seq, and Hi-C sequencing. While SLUR(M)-py generates quality control metrics and useful images/graphs for diagnostic analysis and publication, the pipeline also includes several conversion functionalities; these features allow users to integrate this pipeline into current lab protocols and leverage the outputs from SLUR(M)-py for further analysis.

Here, we demonstrate the utility of our SLUR(M)-py platform in analyzing genomics and epigenomics data using our own ATAC-seq and Hi-C data [40, 41] and Hi-C data from the ENCODE project [42]. We compare the outputs generated by SLUR(M)-py to previous *in silico* efforts as well, including HiCExplorer and Juicer. We show that analysis with SLUR(M)-py greatly improves computational speeds, outpacing previous informatic analysis. Moreover, the data products produced from SLUR(M)-py qualitatively and quantitatively compare to previous outputs made using the Juicer pipeline. Finally, we show that outputs from SLUR(M)-py allow for granular interrogation of contacts between chromosomes as a function of viral infection. This finer resolution analysis reveals a new finding from this data: viral infection does not significantly impact intra-chromosomal contacts, which was missed using other pipelines. Re-analysis of our data also demonstrates additional tools that are democratized within the SLUR(M)-py pipeline, including inter-chromosomal contact frequency analysis.

## Results

### Pipeline overview

SLUR(M)-py combines the best of currently available pythonic and bioinformatic tools and provides parallelization for quickly aligning and processing sequenced reads for studying chromatin structural dynamics and epigenomic modifications. These tools include bio-Python [43], Pandas and Dask Dataframes [44], MACS2 [45–47], fastp [48], and bwa [49]: all were specifically chosen for their speed, efficiency, community familiarity, and capabilities. We combined these tools to form single pipeline to process whole-genome sequencing, ChIP-seq, ATAC-seq, and Hi-C data types. At the command line, SLUR(M)-py only requires a set of paired-end reads and a path to a reference file (in .fasta format) that has been indexed via bwa (Figure 1A and Figure 2A). Like the Juicer pipeline, SLUR(M)-py encodes submissions of scripts to SLURM to efficiently manage parallelized processes within the pipeline (Table 1), first parallelizing across the input read pairs and then the input genome (Figure 1A). However, SLUR(M)-py includes additional scripts to quickly process information that is complementary to Hi-C, such as ATAC-seq data (Figure 1A, Figure 1B, Supplementary Figure S1). Quality control and diagnostic plots are also automatically generated when ATAC-seq or Hi-C datasets are provided (Figure 3, Supplementary Figure S2).

**Figure 1:**
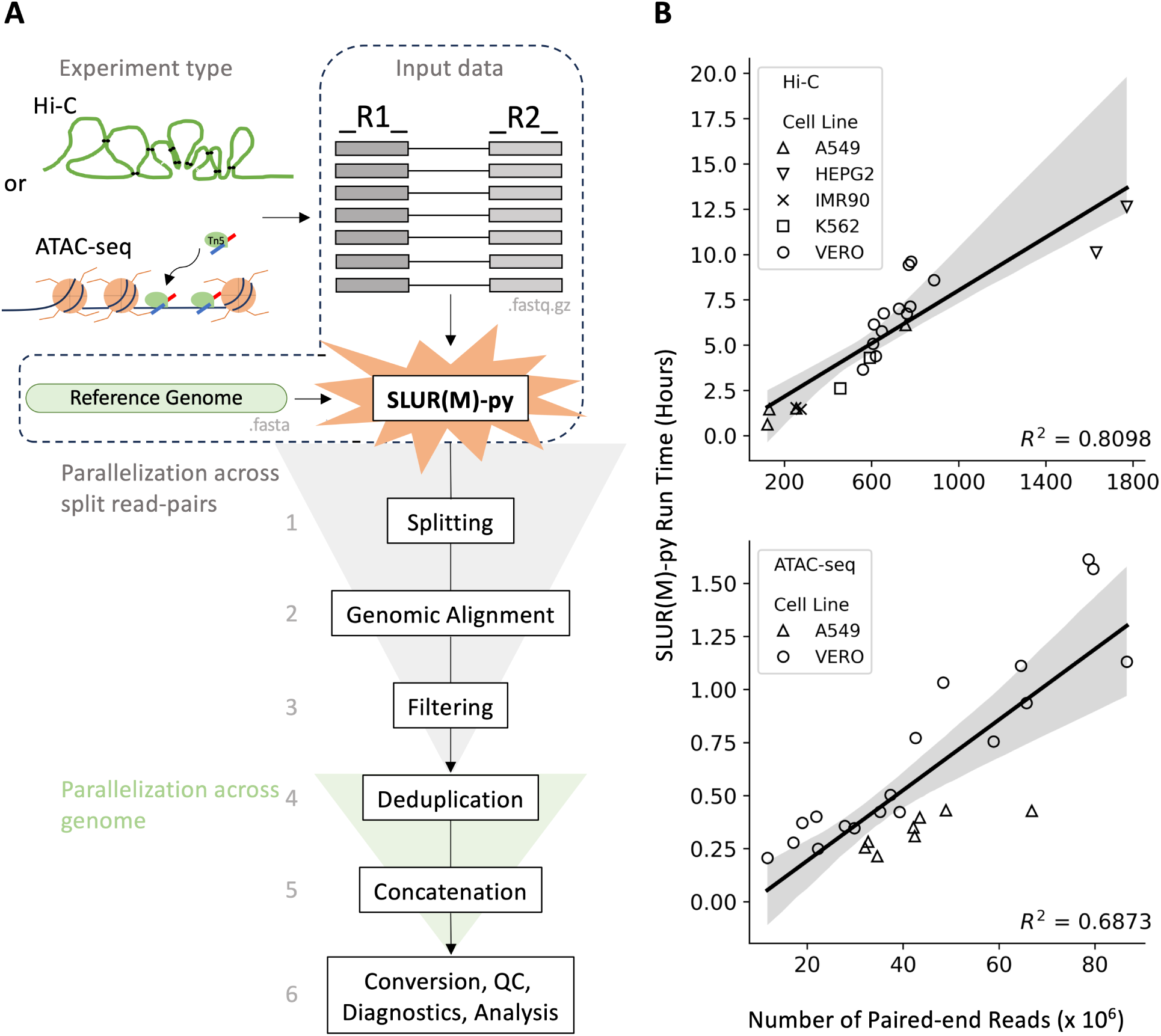
Overview of SLUR(M)-py, a high-performance computing, multi-omic platform for parallelized processing of paired-end reads to study chromatin dynamics. **A**) The basic SLUR(M)-py workflow for processing paired-end sequenced reads from Hi-C or ATAC-seq or experiments. Hi-C measures interacting chromatin in 3D space while ATAC-seq targets open regions of the genome with Tn5, generating paired-end sequencing reads. Paired reads (in .fastq.gz format) and a reference genome (.fasta) are the minimum required inputs to SLUR(M)-py. The SLUR(M)-py pipeline generates jobs and submits them to SLURM which manages and runs tasks for splitting input reads, parallelized genomic alignments, and downstream processes, including filtering, deduplication, and file concatenation. SLURM also sets needed dependencies for file conversion, QC, diagnostics, and further analysis. **B**) The run times (y-axis) of Hi-C and ATAC-seq processing (top and bottom, hours and minutes, respectively) as a function of the number of paired-end reads (x-axis). The cell lines of each sample are depicted as circles (Vero) or x’s, triangles, and squares (human cell lines). Black lines and gray shaded regions represent linear regression models and their 95% confidence intervals (respectively); these models explain 80.98% and 68.73% (*R*^2^) of the variation in run time (top and bottom, respectively).

**Figure 2:**
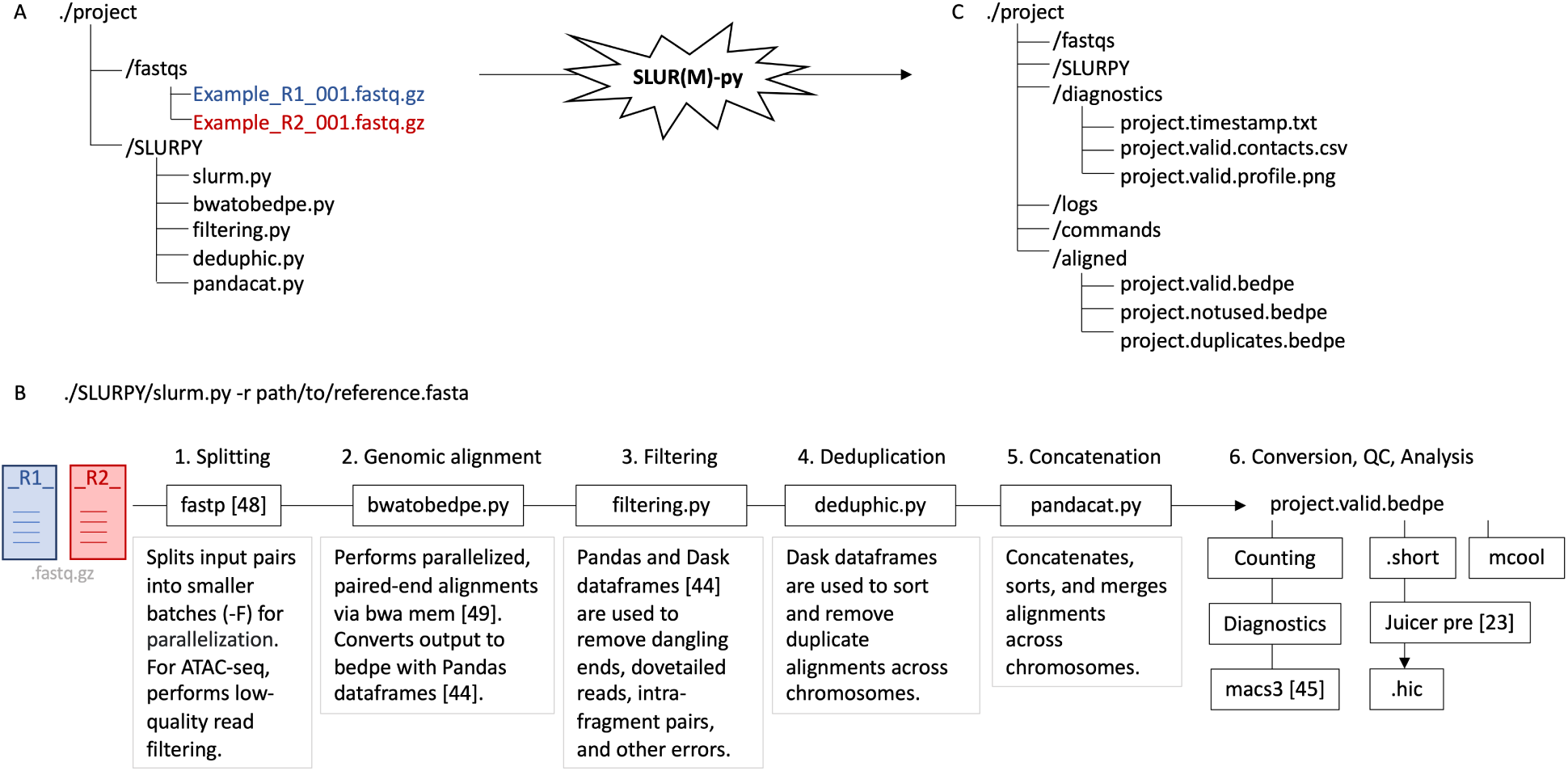
Example project directory and outline of SLUR(M)-py pipeline. **A**) Example project directory structure. Paired fastq.gz files (within the fastqs subdirectory) contain paired-end reads as input into SLUR(M)-py. The directory contains all the scripts and functions associated with SLUR(M)-py and can be soft-linked as a subdirectory. **B**) The processing order of scripts within slurm.py pipeline. To begin, paired fastqs.gz files are split (with fastp) into smaller sets for parallelization. When processing ATAC-seq data, fastp conducts initial quality control, removing low-quality reads, and produces diagnostic reports. Across splits of paired fastqs.gz files, paralleled calls to bwa mem align paired reads to a reference genome (set by the -r argument) are reformatted into bedpe mode (with bwatobedpe.py). A filtering script (filtering.py) segregates alignments into valid and error set pairs. Post alignment and filtering, deduphic.py identifies and removes duplicates, sorting and merging outputs per chromosome. Genome-wide, files are concatenated (with pandacat.py) into final outputs for post processing, including counting, file conversion, and analysis. **C**) After running SLUR(M)-py, diagnostic, logs, and aligned directories are generated containing error logs, quality control plots, and final outputs (respectively).

**Figure 3:**
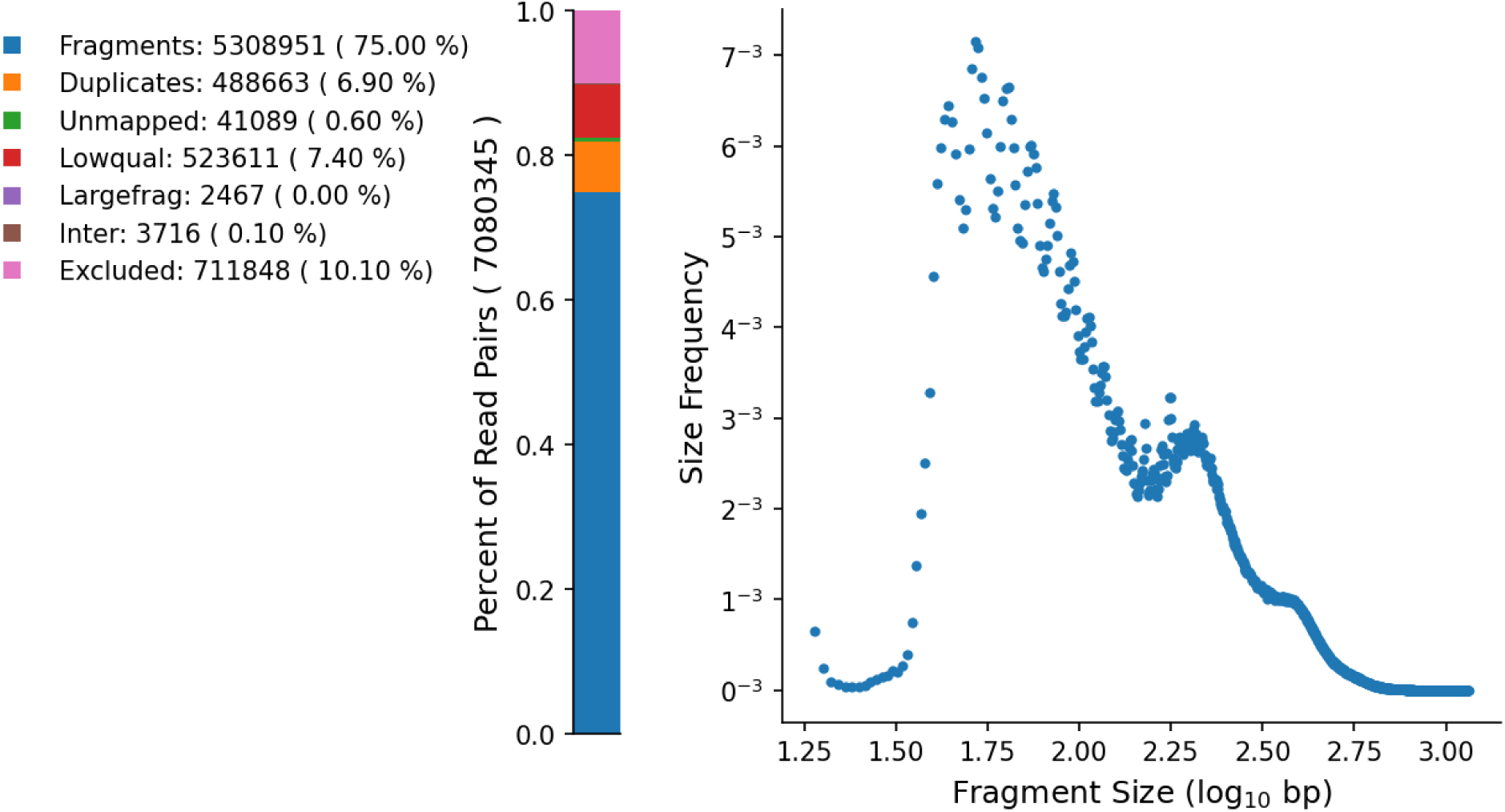
SLUR(M)-py generated diagnostic plot. **Left**) Alignment counts plot (generated by slurm.py) displaying mapping patterns of reads for a Vero ATAC-seq sample. Colors depict different categories of processed reads. **Right**) Fragment size distribution plot autogenerated by SLUR(M)-py pipeline for ATAC-seq analysis. The x-axis denotes the size of fragment formed by aligned, paired-end reads (log_10_) while the y-axis marks the frequency of the fragment size out of the total aligned paired-end reads. Peaks in the curve at approximately 1.75 and 2.25 (along the x-axis) represent numerous fragments with an approximate size 56 and 178 bp respectively.

**Table 1:** Processing features within SLUR(M)-py and published Hi-C pipelines.

|  | SLUR(M)-py | Juicer [23] | HiCExplorer [27] | Fan-c [26] | HiC-Pro [39] |
| --- | --- | --- | --- | --- | --- |
| Multi-omic | Yes | No | No | No | No |
| SLURM management | Yes | Yes | No | No | Yes |
| Pythonic basis & filtering | Yes | No | Yes | Yes | Yes |
| Replicate separating | Yes | No | No | No | Yes |
| Arima compatible | Yes | Yes | Yes | No | Yes |
| Dovetail compatible | Yes | Yes | No | No | Yes |
| Enzymatic libraries<sup>a</sup> | Yes | Yes | Yes | Yes | Yes |
| Calls *bwa mem* with the -5SPM option<sup>b</sup> | Yes | Yes | No | Yes | No |

<sup>a</sup> Restriction site based libraries: MboI, DpnII, Sau3AI, and HindIII
<sup>b</sup>The -5SPM option sets the *bwa mem* command to ignore pairing, skip mate rescue, mark shorter split hits as secondary, and take the alignment with the smallest coordinate as the primary alignment. Often referred to as the "Hi-C option".

The computing environment and dependencies SLUR(M)-py requires are listed on the hosting Github repository and can be easily installed via Python’s package, dependencies, and environment manager, conda [50]. When running SLUR(M)-py on a computing cluster, no specific node type is required. Scripts generated by SLUR(M)-py and submitted to SLURM are node agnostic and do not require a graphical processing unit (gpu) to perform tasks. By default, SLUR(M)-py will instruct SLURM to submit jobs to terabyte (tb) node(s); however, any type or combination of node(s) (or a list of nodes) may be specified with the -P argument in calls to SLUR(M)-py. Alternatively, specific nodes can be listed by name via the --node-list argument. After installation, SLUR(M)-py can be linked within a given project directory (Figure 2A). This project directory must contain a subdirectory named “fastqs” which holds the paired fastq files (in gzipped compressed format [51]) from a sequencing experiment. Multiple replicates (i.e., more than one pair of fastq files) may be held within this directory. Unlike other pipelines (for example see Table 1), the SLUR(M)-py workflow can process multiple replicates (by adding the --nomerge flag at the command line) or merge replicates into a final output throughout processing, which is the default behavior. All of these address challenges we have experienced in analyzing (epi)genomics data sets and exist as improvements to currently available analysis pipelines.

To begin a run, the alignment script within SLUR(M)-py (./SLURPY/slurm.py) requires, 1) a path to a reference file in .fasta format, set by the -r argument, and 2) the presence of gzipped, fastq files (fastq.gz) within the fastqs directory (Figure 2B). By default, SLUR(M)-py expects paired-end reads that represent the sequenced material from Hi-C experiments. This behavior can be modified by passing the additional flags --wgs, --atac-seq or a path to control .bam file (via -c) to slurm.py at the command line, initiating processing of data from whole-genome sequencing, ATAC-seq, or ChIP-seq experiments, respectively (Supplementary Figure S1). A full list of command line options and modifications is listed on the associated SLUR(M)-py github or visible via the help menu (-h). After passing command line arguments, the fastq preprocessing software fastp is used to quickly split input read pairs into smaller sets [48]. The number of paired reads per splits generated by fastp is controlled via the optional “-F” argument. Post splitting of input reads, SLUR(M)-py scripts conduct genomic alignments with the bwa mem algorithm including filtering, deduplication, and concatenation of generated files (Figure 2B); this processing is done in parallel per split. The basic SLUR(M)-py procedure ends with the counting of reads within these files and the removal of temporary files. The final output of this process is a text-based .bedpe file, representing the filtered, paired-end alignments.

In the event of SLURM crashing, node failure, or an error, SLUR(M)-py contains a check point system within the workflow. This feature was included to save data generated at intermittent steps within the pipeline and allow for restarts at specific steps within the protocol. For example, when attempting to process Hi-C sequencing data, after splitting input read pairs with fastp—which is arguably one of the most time-consuming steps within SLUR(M)-py—users may need to restart alignments with bwa mem. This can be done without again splitting input read pairs by passing “-R bwa”, which would direct the slurm.py script to pick up at the step of alignment. In the event of an error, the SLUR(M)-py scripts also include a “checkwork.py” function to identify sources of errors saved within the “logs” directory (Figure 2C). For completely restarting runs, SLUR(M)-py can be passed to a “--restart” flag, should users wish to completely erase, reset, and rerun an attempt. After completing processing with SLUR(M)-py, a “clean” functionality is also included to remove large intermittent and temporary files generated during processing and then compress the final, large text-based files (in .bedpe format) and other outputs with the gzip command.

### Hi-C processing

SLUR(M)-py was initially designed to process sequenced, paired-end reads from Hi-C experiments. Given the underlying chemistry and preparation of Hi-C sequencing, the fragments from these experiments require additional, unique processing (compared to traditional, linear, paired-end reads from whole-genome sequencing) [31]. Briefly, alignments from bwa mem are reformatted at output from the usual sequence alignment maps (.sam format) to a bed, paired-end file (.bedpe); these files are easily passed for further processing to Pandas and Dask Dataframes (Figure 2B). Across splits on input read pairs, alignments are filtered to select valid, informative Hi-C contacts, removing erroneous read pairs representing dangling ends from restriction digestion, pairs that dovetail or map to the same restriction fragment, or present as a self-circle/self-ligation event [31]. After filtering, alignments are segregated by chromosome (taken from the input reference file or a list of chromosomes given by the -G parameter) into separate files. Per chromosomes, paired-end reads are analyzed for duplicate alignments using Pandas and Dask Datarames (Figure 2B). SLUR(M)-py marks and removes duplicate read-pairs that have identical 5’ genomic coordinates and identical sequence signatures, imputed from cigar strings, as previously defined [52]. During this step, entries within each bedpe file representing Hi-C contacts are sorted (from left to right) by genomic position. Post duplicate marking, removal, and sorting, Hi-C contacts are concatenated into a set of final bedpe files within the “aligned” subdirectory, representing valid, unused, and duplicated Hi-C contacts (Figure 2C). This penultimate form of Hi-C data can be converted by SLRU(M)-py into an .mcool file or other file formats for compatibility with Juicer (.short format). Passing a jar file from Juicer tools will que the slurm.py command to auto-generate a .hic file upon completion of Hi-C processing. Our processing of Hi-C data finished in less than 13 hours across all samples reprocessed in this study with average run times of 6.69 and 4.23 hours from samples in Vero and human cell lines, respectively. These run times are faster than our previous run times with current, published pipelines (discussed later in the manuscript).

### ATAC-seq and other processing

SLUR(M)-py is unique among analysis tools in its ability to process other forms of paired-end sequencing data (Table 1). For this purpose, we have encoded the --atac-seq and --wgs flags. Both these flags will configure a run of SLUR(M)-py to process, linear paired-end sequencing data, thereby altering alignment options in *bwa mem*, bypassing checks for Hi-C errors. Furthermore, with the --atac-seq option, output files are converted to be compatible with MACS3 (the most recent version of MACS2 [46]) which is used to identify “peaks” or genomic loci with significant pileup and overlap of mapped sequenced reads. Alternatively, a .bam (or .bedpe) file can be passed to slurm.py at the command line for processing and peak calling of ChIP-seq rather than ATAC-seq data (Supplementary Figure S1).

### Automation of diagnostic plots for ATAC-seq and Hi-C

Upon successful completion of a run, a diagnostics directory is also generated which contains a timestamp (listing the start, end, and total run time) and a visualization (and .csv file) of the read alignment summary. Additionally, for ATAC-seq (and ChIP-seq experiments), read mapping summaries, fragment size distributions (Figure 3) and FRiP scores [18] (Supplementary Table S1) are also automatically calculated and reported to users within the same diagnostics directory. Similarly, for Hi-C analysis, the portion of valid Hi-C contacts (both inter- and intra-chromosomal contacts) and other mapped/processed read pairs are summarized automatically by SLUR(M)-py (Supplementary Figure S2). These diagnostic plots are helpful in determining the success and quality of an experiment.

### Runtime analysis

To test the run time efficiency, flexibility, and fidelity of SLUR(M)-py, we processed both Hi-C and ATAC-seq data from two of our previous studies [40, 41]. Experiments from Venu et al. (2024) in Vero cells include paired ATAC-seq and Hi-C samples taken at 12, 18, and 24 hours post infection with vaccinia virus. In Roth et al. (2023), ATAC-seq assays from A549 cells for several biological replicates (*n* = 8) were collected under nominal laboratory conditions (no perturbation or viral infection). Additional Hi-C data in human cell lines were gathered from the public ENCODE project and repository [42] in the form of raw fastq files and similarly reprocessed using the SLUR(M)-py pipeline. In total, these previously published data sets provided us with twenty-six ATAC-seq (Supplementary Table S1) and twenty-two Hi-C (Supplementary Table S2) samples for processing.

For ATAC-seq samples processed with SLUR(M)-py, the average run times for samples from the A549 simulation study (*n* = 8) and vaccinia virus infection experiments in Vero cells (*n* = 18) were 28 and 40 minutes, respectively, with three datasets taking a little over an hour to completely process the raw paired fastq.gz files into .bedpe files (Supplementary Table S1). Analysis of Hi-C data (*n* = 22) was also relatively quick, finishing in less than 16 hours across all the data analyzed here (Supplementary Table S2). The run times of both Hi-C and ATAC-seq were linear with respect to the number of paired-end reads (Figure 1B, top and bottom, respectively). Indeed, modeling of SLUR(M)-py run time (as a function of counts of input read pairs) explains approximately 80.98% and 68.73% of the variation in run time for Hi-C sequencing and ATAC-seq experiments, respectively (*p*-value *<* 10^−6^, Linear Regression).

### Comparing Hi-C processing

The Juicer pipeline and our SLUR(M)-py Hi-C script both use SLURM to manage jobs and possess Hi-C data; therefore, we wanted to compare our strategy for Hi-C processing against the current standard. After running both the Juicer and SLUR(M)-py Hi-C pipelines on the Vero Hi-C data, we generated .hic files using the processed results from these pipelines and the *pre* command from the suite of Juicer tools [23]. Visually, the Hi-C maps from Vero experiments look similar (Figure 4A, Supplementary Figure S3). For quantitative comparisons, the Hi-C spector reproducibility score [53] was used to quantify the similarity between SLUR(M)-py versus Juicer processed Hi-C maps. Overall, the spector reproducibility scores calculated between Hi-C data processed by SLUR(M)-py and Juicer are biologically reproducible, with a genomic median score ranging from 0.944 to 0.976 across both infected and control samples (*n* = 12). Across these Hi-C samples, most individual chromosomes displayed reproducibility scores higher than 0.80 with the median reproducibility score across samples, per chromosomes all above 0.90 (Figure 4B) signifying high reproducibility between SLUR(M)-py and Juicer pipelines [53].

**Figure 4:**
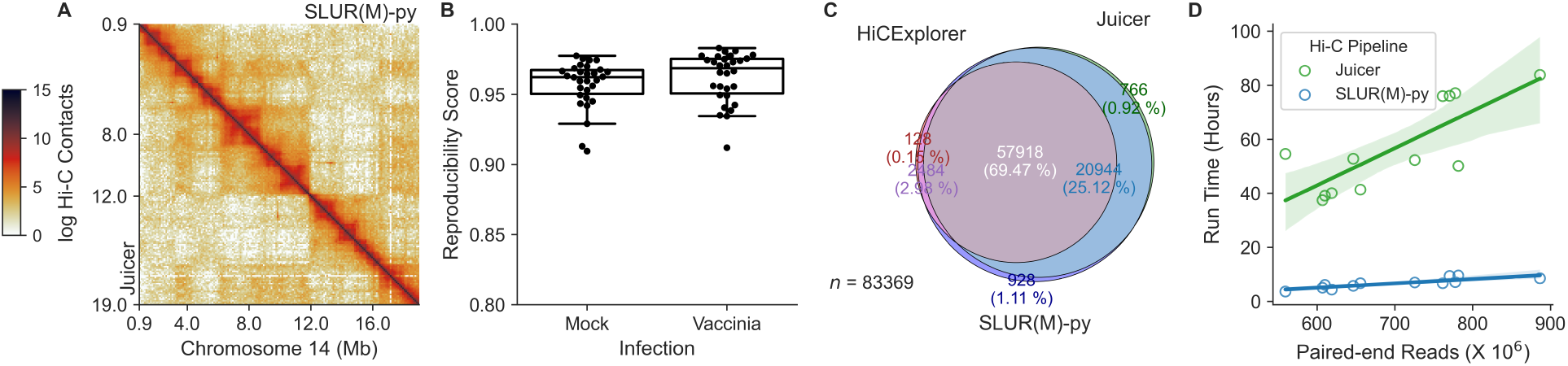
Comparison of Hi-C processing. **A**) Hi-C maps along Vero chromosome 14 from Juicer (lower, left diagonal) and SLUR(M)-py (upper, right diagonal) pipelines. Light to dark orange colors depict increasing Hi-C contact counts (log scale). **B**) Spector reproducibility score between Hi-C maps produced using Juicer versus SLUR(M)-py. The median reproducibility score (y-axis) is shown per chromosome, across Hi-C replicates from Venu et al. (2024) separated by infection (x-axis). Scores above 0.9 are considered highly reproducible [53]. **C**) Overlap of valid Hi-C contacts across pipelines. A random sample of 100,000 read pairs from an A549 Hi-C experiment were processed through HiCExplorer, Juicer, and SLUR(M)-py. A total of 83,369 read pairs were designated as valid contacts by at least one pipeline while 57,918 contacts were designated as valid by all three pipelines. **D**). Distributions of run times (y-axis, hours) as a function of total paired-end reads when processing Vero Hi-C data from Venu et al. (2024) with Juicer (green) pipeline versus SLUR(M)-py (blue). Green and blue lines and shaded regions denote regression models and relating run-time (hours) to total input read pairs.

For two samples (both infected with vaccinia virus), a single chromosome, chromosome 19 (NC_023660.1), displayed scores below 0.80 (Supplementary Figure S4). Examination of read pairs processed by SLUR(M)-py and Juicer revealed that Juicer retains reads that are chimeric singletons (mapping to more than one chromosome in a single read) whose mate goes unmapped, while the scripting in SLUR(M)-py drops a chimeric mapping read(s), with an unmapped mate. This observation and difference in processing, along with viral infection, lead to the low reproducibility scores for this chromosome in these samples. We discuss the detection of ligation events within Hi-C assays between the contig representing the vaccinia virus and the Vero genome later within this manuscript. Overall, visual inspection and quantitative measure of Hi-C maps confirms the high reproducibility between results generated by our SLUR(M)-py pipeline and the Juicer pipeline.

For additional, direct comparison of SLUR(M)-py Hi-C processing to other pipelines, we generated a novel Hi-C data set in the human, A549 cell line using the Arima Hi-C sequencing protocol. Over 700 million paired-end reads were sequenced and processed using our SLUR(M)-py pipeline (Supplementary Table S2). From these data, one-hundred thousand paired-end reads were randomly subsampled and reprocessed using HiCExplorer, Juicer, and SLUR(M)-py. This was done to quickly process reads, mapping them to the human T2T reference genome, and to directly compare their processing (by read name) into Hi-C contacts across all three pipelines. Where possible, similar parameters, like mapping quality scores (*>*= 30), were used in calls for all three pipelines. Across Juicer, HiCExplorer, and SLUR(M)-py, 83,369 out of 100,00 read pairs were retained as valid Hi-C contacts by at least one of the three pipelines (Figure 4C). The percent overlap of read pairs retained as valid Hi-C contacts by all three pipelines (post alignment, processing, and filtering) was 69.47% (Figure 4C). After examining the code-base(s), we postulate that the observed difference in Hi-C contacts is caused by one of the following: 1) differences in mapping approach (specifically options used in the bwa mem aligner, as shown in Table 1), 2) randomness introduced during mapping (specifically the lack of a random seed generator passed to bwa), 3) retention/processing of chimeric singletons (as previously discussed), 4) duplicate marking and handling, and 5) handling of read-pairs mapping to identical DNA restriction fragments (defined by restriction sites and enzymatic digestion). The combined effect of these factors likely leads to large difference in results when the same data is processed by different pipelines and could extend to other Hi-C data processors. This analysis reveals that HiCExplorer had the least number of valid Hi-C contacts (post processing) and the most conservative definition of a “valid” Hi-C contact. By default, SLUR(M)-py retains more read pairs in analysis (like Juicer, those intra-chromosomal contacts that are nearly linear in mapping) compared to HiCExplorer. Thus, we have included an additional preset that may be passed to SLUR(M)-py such that during filtering of Hi-C contacts stricter filtering on alignments and contacts are provided (to better match results from HiCExplorer).

Finally, after compared mappings of Hi-C data (finding broad reproducible with the Juicer pipeline), we tested run times of SLUR(M)-py compared to Juicer. Specifically, we used the early exit before .hic file creation in Juicer and the Hi-C data from the Vero experiments from Venu et al. 2024. Across samples, all the runs with the Juicer pipeline took longer than 24 hours to converge to a pairs-txt file, while all the runs automated with SLUR(M)-py finished in less than 13 hours (Figure 4D). We attribute this increase in computational speed and reduction in run time when using SLUR(M)-py to using fast software for read splitting (fastp [48]), Pandas and Dask Dataframes [44]), and a larger set of initial splits made on input read pairs, which decreased overall memory requirements and increased the number of parallel processes.

### Duplicate marking

During initial library preparation, PCR amplification can lead to the propagation of duplicate sequenced fragments. Alternatively, a single amplification cluster can be incorrectly identified as multiple clusters by the optical sensor, leading to the creation of optical duplicates. Most current practices suggest that duplicate fragments should be identified and removed to eliminate bias and the inclusion of artifacts in downstream analyses [18, 22, 23, 40, 54, 55]. However, for some omics analyses, marking duplicates may be unwarranted. For example, during genetic variant calling from whole genome sequencing, a recent study has suggested that marking and removing duplicates is memory intensive and unnecessary [56]. Similarly, another study examining duplicates within ATAC-seq data discovered that many duplicate fragments (identified by position) are not always the result of PCR amplification, but rather, true sequenced fragments created by the same, small open regions being sequenced multiple times [57]. For identifying duplicate alignments, SLUR(M)-py filters alignments using the 5’ genomic positions of fragments, strand orientation, and sequence signal/nucleotide variation; matching the definition of duplicate previously defined by other tools like Picard and Samblaster [52]. Given that users may wish to skip this step, duplicate marking and removal is optional within SLUR(M)-py, controlled by the addition of the --skipdedup flag at the command line.

To test the effect of skipping duplicate markings and removal on processing times and results generated from ATAC-seq data, we ran several timing tests using the ATAC-seq data (in A549 cells) generated in Roth et al. (2023). Skipping duplicate marking and removal had no significant impact on overall run times (Supplementary Figure S5A). While we did see a slight increase in processing time—an insignificant 1.18 minutes on average (*p*-value *>* 0.05, Wilcoxon signed-rank test)—this was attributed to an increased data size processed by penultimate steps in the pipeline (such as MACS3) and not duplicate marking and removal. Similarly, while we saw small increases in the number of valid ATAC-seq fragment counts (Supplementary Figure S5B) and FRiP scores (Supplementary Figure S5C) post processing (when retaining duplicates), neither of these increases were deemed significant (*p*-value *>* 0.05, Wilcoxon signed-rank test).

While marking and removing duplicates has little effect on processing times and diagnostics of these experiments, the overall effect on detected, significant peaks (from MACS3) was unclear. In most samples, the number of peaks detected by MACS3 increased when duplicate read pairs were retained (Supplementary Figure S5D), adding an average (median) of nearly 6,371 peaks to the overall peak counts across experiments in the A549 cell line. Yet in two (of the eight) A549 samples, fewer peaks were identified when retaining duplicates during processing (Supplementary Figure S5D). We hypothesize that in these two, outlying samples, duplicate reads (when retained in the analysis) increased the level of background noise, lowering the overall number of peaks deemed significant by MACS3. A note of caution to users, for ATAC-seq experiments, excluding duplicate marking and removal may not affect the total processing time, but it is likely to alter the number of significant peaks identified via peak callers (like MACS3).

### Inter-chromosomal interaction measurements

Within the cellular nucleus, chromosomes segregate into territories [58]. Consequently, most sequenced, paired-end reads from Hi-C experiments will map within a given chromosome and are labeled as intra-chromosomal contacts [31]. However, a small portion of read pairs map between chromosomes, representing interactions between chromatin from different, neighboring chromosomes; these valid Hi-C contacts are referred to as inter-chromosomal contacts. Traditionally, the ratio of inter-to intra-chromosomal contacts is monitored by studies but methods of scoring inter-chromosomal contacts have not been developed [59].

To broadly quantify the interaction between chromosomes, SLUR(M)-py provides an estimate of the inter-chromosomal contact scores [15, 60]. The inter-chromosomal contact score, as described in Lieberman-Aiden et al. (2009) and Duan et al. (2010), is a broad quantification of the interaction frequency between two chromosomes. Specifically, this score is a ratio calculated by dividing the number of observed contacts between two chromosomes by the expected number of contacts between them, given the total number of inter-chromosomal contacts involving said chromosomes. For visualization of these scores, SLUR(M)-py generates a heatmap using the Seaborn plotting package in python (Figure 5A) [61]. These calculations are also included within the diagnostic plot when processing Hi-C data (Supplementary Figure S2).

**Figure 5:**
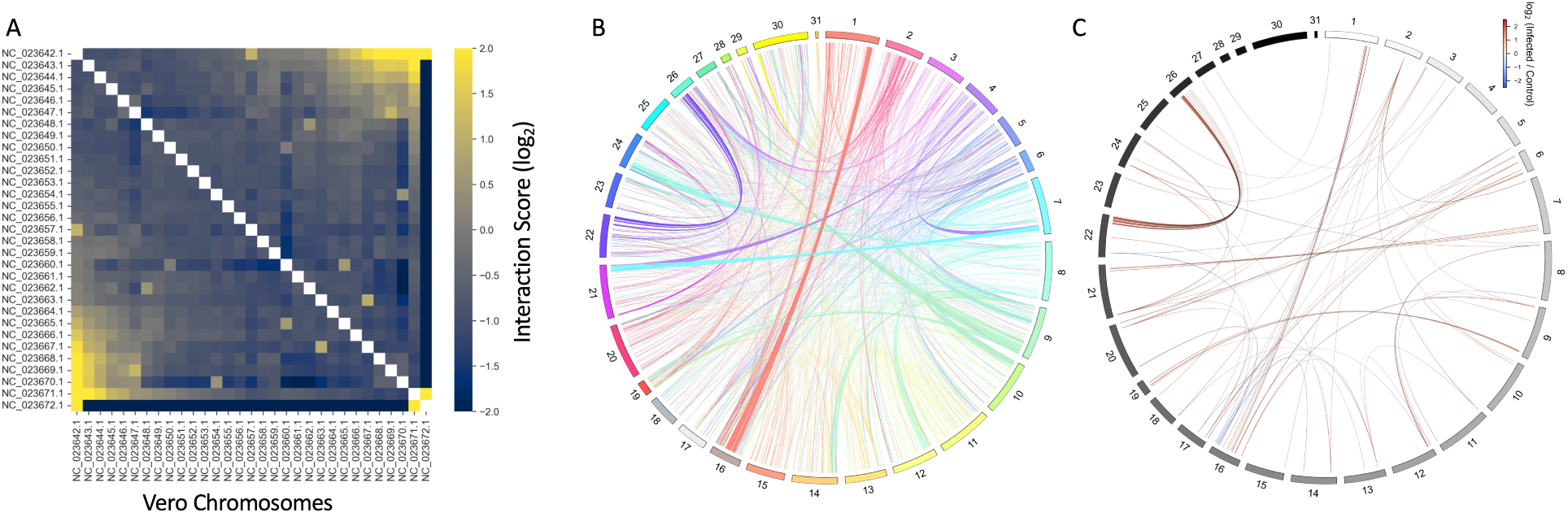
Inter-chromosomal analysis automated within the SLUR(M)-py pipeline. **A**) The interaction scores (log_2_) between chromosomes (x- and y-axis,) are measured as the ratio of contacts between a pair of chromosomes and the expected number of inter-chromosomal contacts of those chromosomes. **B**) Significant inter-chromosomal interactions between Vero chromosomes. Colored arcs represent 500 kb regions with a significant number of Hi-C contacts between chromosomes as measured via a genome-wide binomial model. **C**) The fold-change of inter-chromosomal interactions with significant differences between control Hi-C map and the Hi-C map from Vero cells post 24 hours of exposure to the vaccinia virus.

When examining the inter-chromosomal contacts within Hi-C data from Venu et al. (2024), we detected an artifact of the ligation events between the vaccinia virus and the Green Monkey reference genome. The Green Monkey genome as reference is much more complete and less complicated (with only 31 contigs/chromosomes) compared to the published Vero genome, which requires significant improvement [62]. Designed with host-pathogen infection experiments in mind, SLUR(M)-py can filter out alignments mapping to pathogen contigs, assuming the pathogen contig has been added as an additional contig within the reference genome. These artifacts were present in both the Hi-C and ATAC-seq data and the number of detected vaccinia virus fragments was proportional to the number of sequenced reads (*p*-value *<* 0.05, *R*^2^ = 0.923, Supplementary Figure S6). Within the Hi-C experiments, these fragments make up a small portion (less than one percent) of the ligation events and only appeared in assays from infected cells (Supplementary Figure S7). Read pairs mapping to the vaccinia virus contig are recoverable from ATAC-seq experiments. However, no read pairs were identified that span vaccinia virus and genome in this assay.

The apparent ligation between chromosomes and virus within the Hi-C experiment was further explored post processing with SLUR(M)-py. Specifically, we examined the anchoring points of paired reads, isolating pairs with one read mapping to the virus contig and the other belonging to the Green Monkey genome. In depicting the mapping of these coordinates, we discovered that the supposed ligation events between the virus and genome are uniformly distributed (Supplementary Figure S8). Furthermore, across chromosomes, the proportions of paired reads connecting the virus and genome are linear with respect to the size of the chromosome (*p*-value *<* 2_10_, *R*^2^ = 0.92, Supplementary Figure S9). These observations, along with historic, experimental evidence that vaccinia virus does not enter the nucleus, led us to conclude that these observed ligation events are false positive, inter-chromosomal contacts. From these events, we can predict a lower bound for the number of false ligation events in our experiments, making up approximately 0.345 percent of observed inter-chromosomal ligations (Supplementary Figure S10A). Furthermore, these false positive events were independent from the number of sequenced reads within Hi-C experiments (*p*-value *>* 0.05, linear regression, Supplementary Figure S10B).

Broadly, in the Vero experiments from Venu et al., the number of inter-chromosomal contacts was approximately 10% (of the total set of valid Hi-C contacts) within each Hi-C experiment (Supplementary Figure S7). This overall proportion of inter-chromosomal contacts did not significantly vary between controls and virus-infected samples. Previously, we had observed alterations in chromatin interaction frequencies between control and infected samples [41]. However, this analysis was only conducted at the scale of chromosomes. Here we performed additional, deeper analysis of the interactions captured between chromosomes using a combination of the output from our SLUR(M)-py, pipeline and PyDESeq2 [63]. The contacts between chromosomes were quantified by generating contiguous bins, 500 kb in size, left to right, along each chromosome across samples, generating 841,610 unique, inter-chromosomal bins (Figure 5B). The number of contacts between pairs of chromosomes were counted per bin and used as inputs into PyDESeq2 to search for differentially altered inter-chromosomal contacts due to vaccinia virus infection.

Collapsing across time, we observed poor separation of infected and control samples when performing principal component analysis on the binned inter-chromosomal counts (Supplementary Figure S11A, left). Similarly, across the infection time-course only 66 of 841,610 bins (less than one percent) formed between interacting chromosomes were identified as significantly differential (adjusted *p*-value *<* 0.001, absolute fold-change *>* 1, Supplemental Figure S11A, right). Performing linear contrasts and examining differential inter-chromosomal contacts between chromosomes at each hour post infection (hpi), we saw no effect at 12 hpi (Supplementary Figure S11B), a minimal effect at 18 hpi (Supplementary Figure S11C), and the greatest separation of infected and control samples at 24 hpi (Supplementary Figure S11D). At this final time point (24 hpi), we identified only a small number of inter-chromosomal regions (13 and 138) with significant enrichment in fold-change, with respect to controls (negative fold-change) or infected (positive fold-change) samples (respectively) between chromosomes (*p*-value *<* 0.001, absolute fold-change *>* 1, Figure 5C, Supplementary Figure S11D). This result, the identification of only a small number of differential inter-chromosomal regions, suggests that changes in contacts between chromosomes are not a hallmark of vaccinia virus infection.

## Conclusions and Future Directions

SLUR(M)-py is a SLURM powered, pythonic pipeline for performing parallel processing of paired-end sequencing profiles prepared from epigenomic and genomic experiments. SLUR(M)-py contains protocols and code written in python for analyzing DNA sequences from many of the standard assays used within epigenomic studies. These pythonic scripts allow users to process paired-end sequenced reads from whole-genome, Hi-C, ATAC-seq, and ChIP-seq assays using just SLUR(M)-py. In this regard, SLUR(M)-py is a multi-omic pipeline, designed to run within a HPC, that eliminates the need for researchers to install and maintain multiple pipelines.

Underlying SLUR(M)-py is unique Python code that interacts with the simple Linux utility for resource management system, SLURM. With SLURM, one can parallelize, stack, and schedule dependencies necessary for end-to-end omics analysis. Unlike currently published pipelines, SLUR(M)-py is not dependent upon a gpu node for processing. SLUR(M)-py can schedule jobs to run on any node type integrated within a system managed by SLURM. By default, runs of SLUR(M)-py will seek a tb node. All the results presented within this paper were generated using tb nodes. Our experience has shown that jobs submitted to gpu nodes tend to complete faster, which is expected given the difference between tb and gpu nodes. The flexibility of node choice allows users to run multiple submissions of SLUR(M)-py across several nodes, which is especially beneficial for high performance computing environments with a heterogeneous node architecture.

We developed SLUR(M)-py using the sequencing data from Hi-C and ATAC-seq assays collected from mammalian cells during viral infection experiments [41]. Unlike current high-performance computing bioinformatics pipelines, the names of the contig representing the viral vector, mitochondrial DNA, or sex chromosomes (chrX and chrY for example) can be passed to SLUR(M)-py as additional inputs. This feature allows users to monitor the level of contamination from these sources or to mask their alignments, which are removed early on in processing, saving computational time and cost. Conversely, a list of select contigs from within a reference genome can be passed to SLUR(M)-py to limit analysis to those chromosomes of interest. Furthermore, these features provide flexibility to the SLUR(M)-py workflows which are not restricted to just human systems and cell lines with expected chromosome names such as chr1, chr2, and chrM, representing the first autosomes and the mitochondrial contig (just to name a few examples). Indeed, SLUR(M)-py is designed to be species agnostic. As an example of this, with SLUR(M)-py both Hi-C and ATAC-seq data were reprocessed from human cell lines (taken from the ENCODE consortium) and the African Green Monkey (*Chlorocebus sabaeus*), Vero cell line, produced by Venu et al. (2024). In both cases SLUR(M)-py auto-generated the metrics for quality control, produced the expected data packages for further analysis, and rendered figures for publication without error.

SLUR(M)-py demonstrates quick run-times relative to other pipelines. To our knowledge, Juicer and HiC-Pro are the only other Hi-C workflows that program and submit tasks to SLURM. For comparison, we reanalyzed our previously published Hi-C data [41], which was initially processed using the Juicer pipeline. The run times of SLUR(M)-py were faster than runs with the Juicer pipeline when reprocessing our data from viral infection experiments in the Vero cell line. Additionally, across the genome, the Hi-C maps generated with the outputs from the SLUR(M)-py pipeline match those Hi-C maps made using the Juicer pipeline with high reproducibility. However, SLUR(M)-py has the advantage over Juicer of being able to quickly and effectively analyze multiple types of epigenomics data sets, making bioinformatics analysis easier and more efficient for the end user.

Using SLUR(M)-py we set out to explore the interactions between chromosomes within the Vero cell line. SLUR(M)-py provides scripts for calculating the interaction between chromosomes, and much like human cell lines [15], we see that smaller chromosomes interact more often with each other than with larger chromosomes in C. sabaeus. We expanded on this analysis along chromosomes (across 500 kb chromosome bins) to identify regions with differential inter-chromosomal contacts due to viral infection. Across the infection time course, this new analysis showed little difference suggesting that intra-chromosomal interactions are more impacted than inter-chromosomal interactions in response to vaccinia virus exposure. The largest difference (*<* 1%) in inter-chromosomal contact frequencies was seen at 24 hours post infection. Thus, unlike the dynamics in loop structures (due to infection) observed along chromosomes [41], the changes in contacts between chromosomes across time are not associated with viral infection. For other studies seeking to explore inter-chromosomal dynamics and their role in responding to viral infection, or other environmental perturbations, we have generated a set of scripts within SLUR(M)-py to aid in this analysis.

When analyzing contacts between chromosomes, we utilized SLUR(M)-py to explore artifacts that appear in those experiments from Venu et al. (2024). Specifically, we identified several ligation events between the contig representing the vaccinia virus and the chromosomes of the Green Monkey genome. The DNA of the vaccinia virus does not translocate into the nucleus, and therefore, its DNA does not directly interact with the host chromosomes [64–66]. It follows that crosslinking between the host and vaccinia virus genomes after formaldehyde exposure during Hi-C library preparation is impossible. Thus, we determined these events (which are rare) are due to ligation errors produced after crosslinking. We estimated the rate of these false ligation events that occur within Hi-C experiments, is less than one percent. This finding should provide investigators with higher confidence that inter-chromosomal interactions are not dominated by random noise. For viral infection studies, SLUR(M)-py scripts have the option to remove these false contacts.

For the ATAC-seq data analyzed here—with an average of 21 million paired-end reads—the average run times were a little over half an hour (36.4 minutes). With these data and short runtimes, we investigated the effect on the average runtime when skipping duplicate marking and removal. The effects on runtimes were minimal, and skipping duplicate marking (and removal) is estimated to save users, on average, a little over five minutes. However, skipping duplicate filtering led to an increased count of significant loci detected within each ATAC-seq sample (an additional six-thousand peaks). It has been demonstrated that the nature of ATAC-seq leads to an overestimate of duplicate read pairs [57]. Briefly, small accessible regions create a bias in the final set of sequenced regions. While these regions are biologically reproducible, marking duplicates simply by removing fragments with duplicate positions (the traditional method) affects the total end counts of detected significant loci in ATAC-seq data [57]. Currently, the SLUR(M)-py pipeline is unable to retain duplicate read-pairs based on position and is only able to remove them. Future applications will try to estimate those read pairs that are marked as duplicates yet are biologically valid and worthy of retention. Thus, retaining or removing duplicates within a run of SLUR(M)-py we leave up to the users, controlled by the inclusion of the “--skipdedup” flag at the command line.

In closing, SLUR(M)-py is designed to be a flexible pipeline for quickly processing data from epigenomic experiments. Additional tools and features are currently under development. For example, future versions of SLUR(M)-py will be able to process long read sequences from Oxford Nanopore or PacBio sequencing [67,68]. Protocols for processing sequencing data like RNA-seq, Cut&TAG sequencing [69–71], and genetic variant detection are also under development. While not explicitly tested, SLUR(M)-py should be able to process data from single cell experiments too, assuming the input data is in the form of paired-end sequences. SLUR(M)-py is publicly available and hosted on Github and able to be copied, forked, or modified by the scientific community. It is our hope that SLUR(M)-py contributes towards epigenomic and 3D genomic studies of chromatin conformation and architecture.

## Methods

### Data included, preparation, and processing

The raw data from ATAC-seq and Hi-C sequencing experiments generated from Venu et al. (2024) and used in this study are available in the SRA repository, under the BioProject accession number PRJNA1037174. For details on the viral infection experiments in Vero cells infected with the Vaccinia virus see Venu et al. (2024). Using the SLUR(M)-py, pipeline the paired-end reads from ATAC-seq and Hi-C experiments were aligned to the Green Monkey (*Chlorocebus sabaeus*) reference genome from NCBI, named GCF_000409795.2_Chlorocebus_sabeus_1.1_genomic.fasta [41]. The sequence of the Vaccinia virus (accession number U94848) was downloaded from GenBank and added to this reference genome as an additional contig. This reference file (in fasta file format) was then indexed using bwa [49]. For the Vero data sets (both ATAC-seq and Hi-C) the call to the slurm.py script were modified (using the -G parameter) to include only the 29 fully assembled autosomes (Chromosomes 1 through 29) and the two sex chromosomes (X and Y) with NCBI contig names NC_023642.1 – NC_023672.1. The calls to slurpy.py were also modified to include the removal of reads mapping to the Green Monkey mitochondrial contig (-M NC_008066.1). For analysis of ATAC-seq samples from the Vero cell line, the flag --atac-seq was added to calls of slurm.py. Apart from the inclusion of the -G and -M parameters, all other default settings in SLUR(M)-py scripts were utilized for analysis of the Vero samples.

Details on the creation, handling, and sequencing of ATAC-seq experiments using the A549 cell line presented here in timing and duplicate marking analysis are included within Roth et al. (2023). Briefly, raw, paired-end sequenced reads from eight biological replicates were downloaded from the sequence read archive from the bioproject PRJNA975595 with accession numbers SRR24717527–SRR24717534. Using SLUR(M)-py with the --atac-seq flag and default settings, read pairs per sample were aligned to the telomere-to-telomere human reference genome, version 2 [72].

As part of this study, a Hi-C library was also prepared (using the Arima Hi-C library kit) from a sample of A549 cells. The methods for cellular culturing, handling, library preparation, and sequencing are exactly similar to those described previously in Venu et al. (2024), however using A549 cells rather than Vero cells. Sequenced pair-end reads were also mapped to the telomere-to-telomere human reference genome, version 2 [72]. Raw fastq files from this sample are listed under the SRA accession number XXXXXXXXX. For comparison of Hi-C processing software, 100,000 random paired-end reads were selected from this sample for processing.

### Other data used

Nine sets of paired-end sequences representing Hi-C data collected on human cell lines were taken from the ENCODE project and data repository [22, 42]. These included experiments conducted within the A549, HepG2, IMR90, and K562 cell lines. Please refer to Supplementary Table S2 for the list of ENCODE accession numbers for each sample. These data were each aligned to the telomere-to-telomere human reference genome, version 2 [72] using SLUR(M)-py with default settings.

### Hi-C analysis and comparison

For comparative analysis, the Hi-C samples from experiments within the Vero cell line were aligned to the *C. sabaeus* reference genome modified with the addition of the vaccinia contig (as described above) using the Juicer pipeline [23]. For processing of each replicate with Juicer, the early exit parameter (-E) was also added to end the processing of Hi-C contacts after the creation of the merged, sorted no duplicates text file but prior to the creation of an .hic file. Post processing, .hic files were generated with the pre command from Juicer tools (version = 1.22.01), using both the text-based outputs from Juicer and SLUR(M)-py workflows as inputs. Across both pipeline, a mapping quality score of thirty was used to filter alignments. To compare the .hic files made using the Juicer pipeline to those made using SLUR(M)-py, the spector reproducibility score [53] was calculated for the main autosomes and sex chromosomes at 500 kb resolution across the sampled Hi-C maps (*n* = 12). Runs of SLUR(M)-py auto-generate timestamps and calculate total run time(s). For runs with the Juicer pipeline, we estimated the total run time from start and end timestamps of logs within the Juicer debug output directory. These run times were brought into python for comparison, analysis, and plotting.

### Testing of ATAC-seq analysis

For the analysis of processing times when analyzing ATAC-seq and Hi-C data using SLUR(M)-py, the auto-generated timestamps across runs were transformed into total minutes and hours then regressed against the number of input, paired-end reads using a linear model. For testing the effect of duplicate marking and removal on overall runtimes and processing of ATAC-seq samples, the slurm.py script within SLUR(M)-py was re-run twice for each sample, with and without marking duplicates, post alignment with bwa mem by passing the --skip-dedup flag to the slurm.py. Autogenerated time stamps from these runs were brought into Python for calculating the change in processing time. The auto generated diagnostic files, which contain information on the number of peaks and summits detected with MACS3, for each run were similarly analyzed.

### Inter-chromosomal analysis

Using the SLUR(M)-py script, the number of Hi-C contacts between chromosomes were counted and used to calculate the inter-chromosomal interaction score (IS) following methods from and Lieberman-Aiden et al. (2009) and Duan et al. (2010). Briefly this is calculated for a pair of chromosomes *i* and *j* using the following formula: IS(*i, j*) = *N_ij_/*((*N_i_/N*)(*N_j_/N*)(*N*)). Where *N_i_* and *N_j_* are the number of inter-chromosomal contacts involving chromosomes *i* and *j* (respectively), *N_ij_* is the number of inter-chromosomal contacts between chromosomes *i* and *j*, and *N* is the total number of inter-chromosomal contacts [15, 60]. These scores were visualized using Seaborn’s heatmap function in Python on a log_2_ scale [61].

For visualization of inter-chromosomal contacts, contacts between chromosomes were counted every 500 kb. These inter-chromosomal bins were used in a binomial model from Kaufmann et al. (2015) to identify significant interactions every 500 kb between chromosomes [73]. The Circos, circle plot library [74] in Python was used to plot significant 500 kb bins (binomial test, *p*-value *>* 0.001) between chromosomes.

For differential, inter-chromosomal analysis of Vaccinia infected Vero samples, the inter-chromosomal contacts every 500 kb were counted across replicates and samples and compared between vaccinia infected and control Hi-C maps at 12, 18, and 24 hours post infection. Specifically, the counts of binned inter-chromosomal contacts were used as input into PyDESeq2 [63]. Linear contrasts were performed at each time-point post infection. An adjusted, *p*-value threshold of *p*-value *<* 0.001 and absolute fold-change greater than one were used to establish significant differential contacts between chromosomes.

## Software availability

The code base for SLUR(M)-py is currently under development and hosted on Github at: https://github.com/SLUR-m-Py/SLURPY

## Acknowledgments

We would like to thank members of the Genomics and Bioanalytics Group and the Biosystems Administrators at Los Alamos National Laboratory for their helpful comments on this manuscript. We would also like to thank members of the Genomics Sequencing Core at Los Alamos National Laboratory for sequencing of genomic data presented in this manuscript. We would also like to acknowledge and thank the ENCODE Consortium and the members of the Dekker lab of University of Massachusetts, the Aiden-Liebermann lab of Baylor University, and the Reddy lab of Duke University for generating the Hi-C data sets used within this study. This material is based upon work supported by the U.S. Department of Energy, Office of Science, through the Biological and Environmental Research (BER) and the Advanced Scientific Computing Research (ASCR) programs under contract number 89233218CNA000001 to Los Alamos National Laboratory (Triad National Security, LLC) awarded to CRS and SRS and supported by the Los Alamos National Laboratory Directed Research Grant (20210082DR) awarded to SRS.

## Author contributions

CR develops and maintains SLUR(M)-py and its associated software. VV tested, proposed user features, provided feedback, and assisted in development of SLUR(M)-py software. VV conducted cell culture and library preparation for Hi-C sequencing. CR and VV interpreted and summarized Hi-C results. SB added code, performed debugging, ran testing, and made edits to the SLUR(M)-py pipeline and documentation. CR performed the analysis and generated figures. CRS and SS provided funding and materials for the research presented within this paper. CR and CRS wrote the manuscript. All authors provided edits and comments on the manuscript.

---

**Figure S1:**
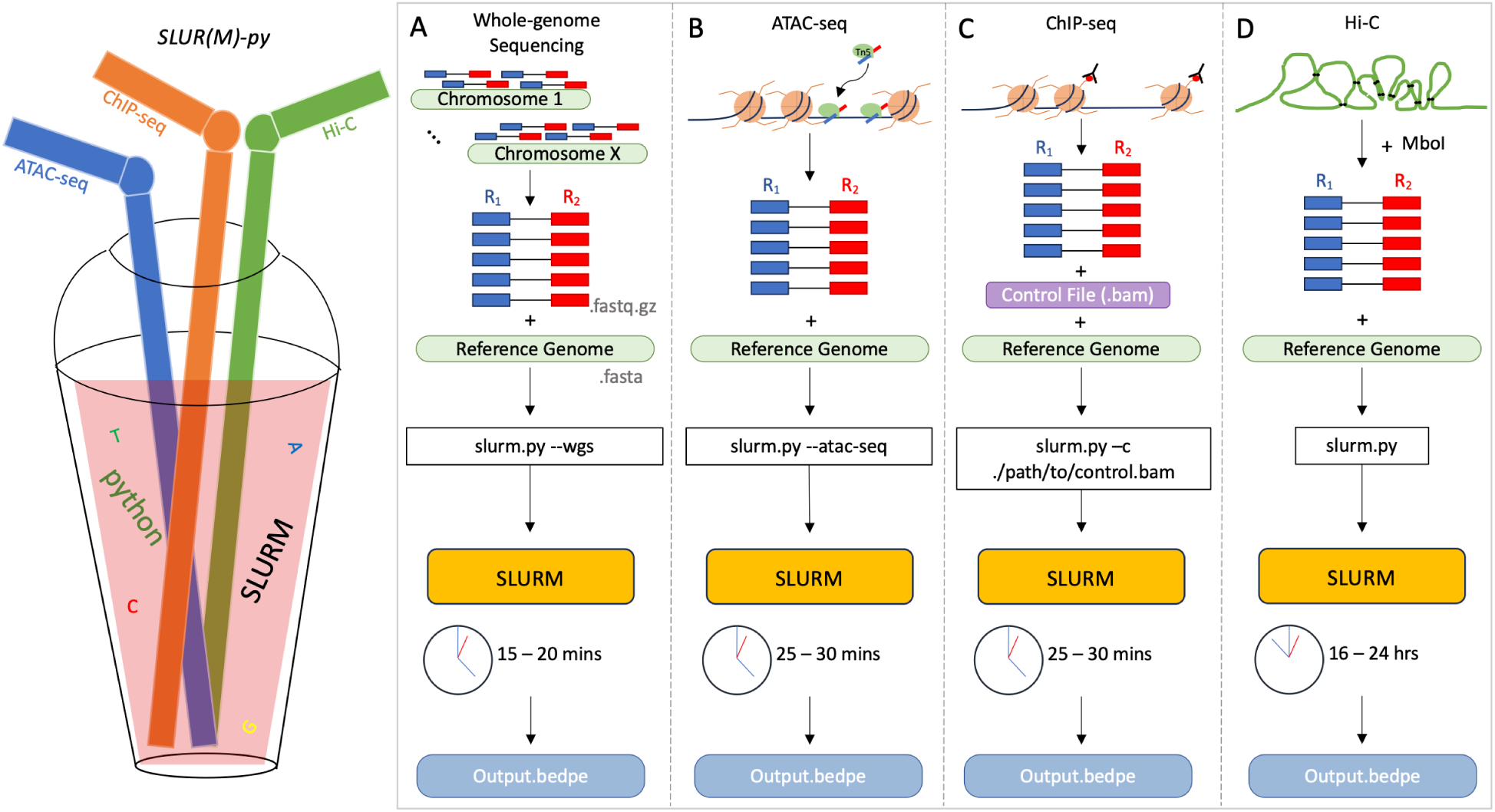
Overview of SLUR(M)-py; a high-performance computing, multi-omic platform for parallelized processing of paired-end reads to study chromatin dynamics. **A**) The basic SLUR(M)-py workflow for processing paired-end, whole-genome sequences. **B**) ATAC-seq targets open regions of the genome with Tn5, generating paired-end sequencing reads. These reads (in .fastq.gz format) and a reference genome (.fasta) are the minimum requirements passed to slurm.py in ATAC-seq mode (addition of --atac-seq flag). From slurm.py, jobs are passed to SLURM to parallelize alignments, set dependencies, and que downstream processes, quickly generating output in bedpe format. **C**) ChIP-seq experiments target modified histone tails to generate paired-end reads after sequencing. For processing ChIP-seq experiments, the slurm.py script requires the addition of a control file (in .bam or .bedpe format). **D**) Overview of inputs for the Hi-C processing. Paired-end reads from Hi-C experiments, along with the name of the fragmentation enzyme, and the reference genome are passed to slurm.py as inputs.

**Figure S2:**
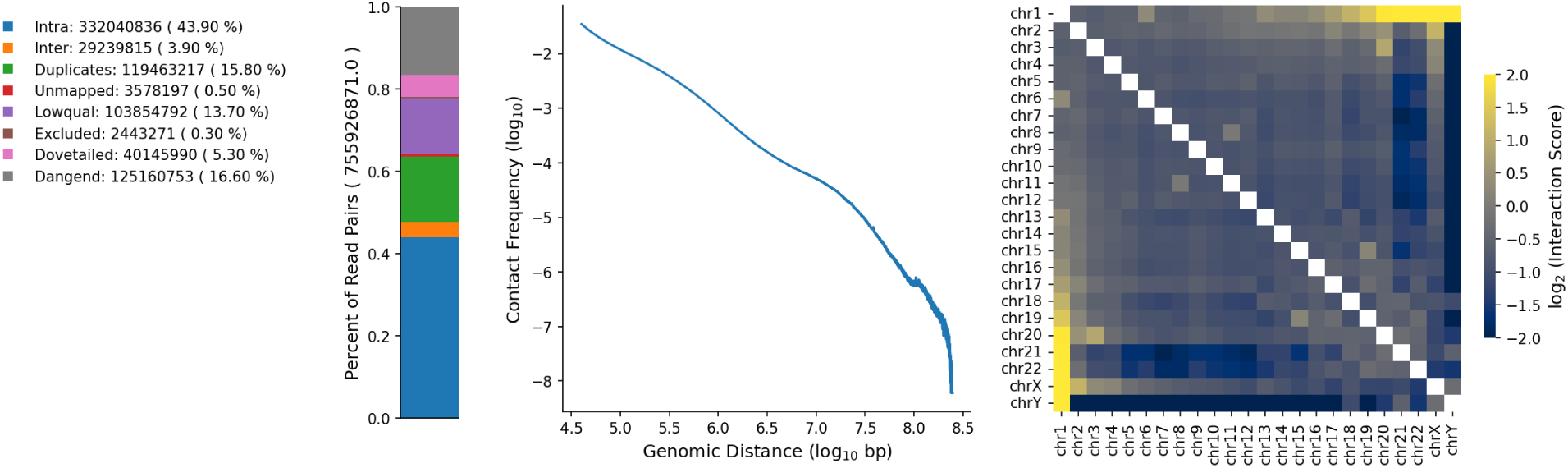
Hi-C diagnostic plot generated by SLUR(M)-py. **Left**: A portion plot showing the portion of paired-end reads kept in analysis as valid Hi-C contacts (blue and orange colors) and those removed during processing. **Middle**: A distance decay profile calculated from intra-chromosomal Hi-C contacts. **Right**: The inter-chromosomal contact score calculated between chromosomes from valid Hi-C contacts. Yellow colors indicate chromosomes with stronger interactions compared to others (blue colors).

**Figure S3:**
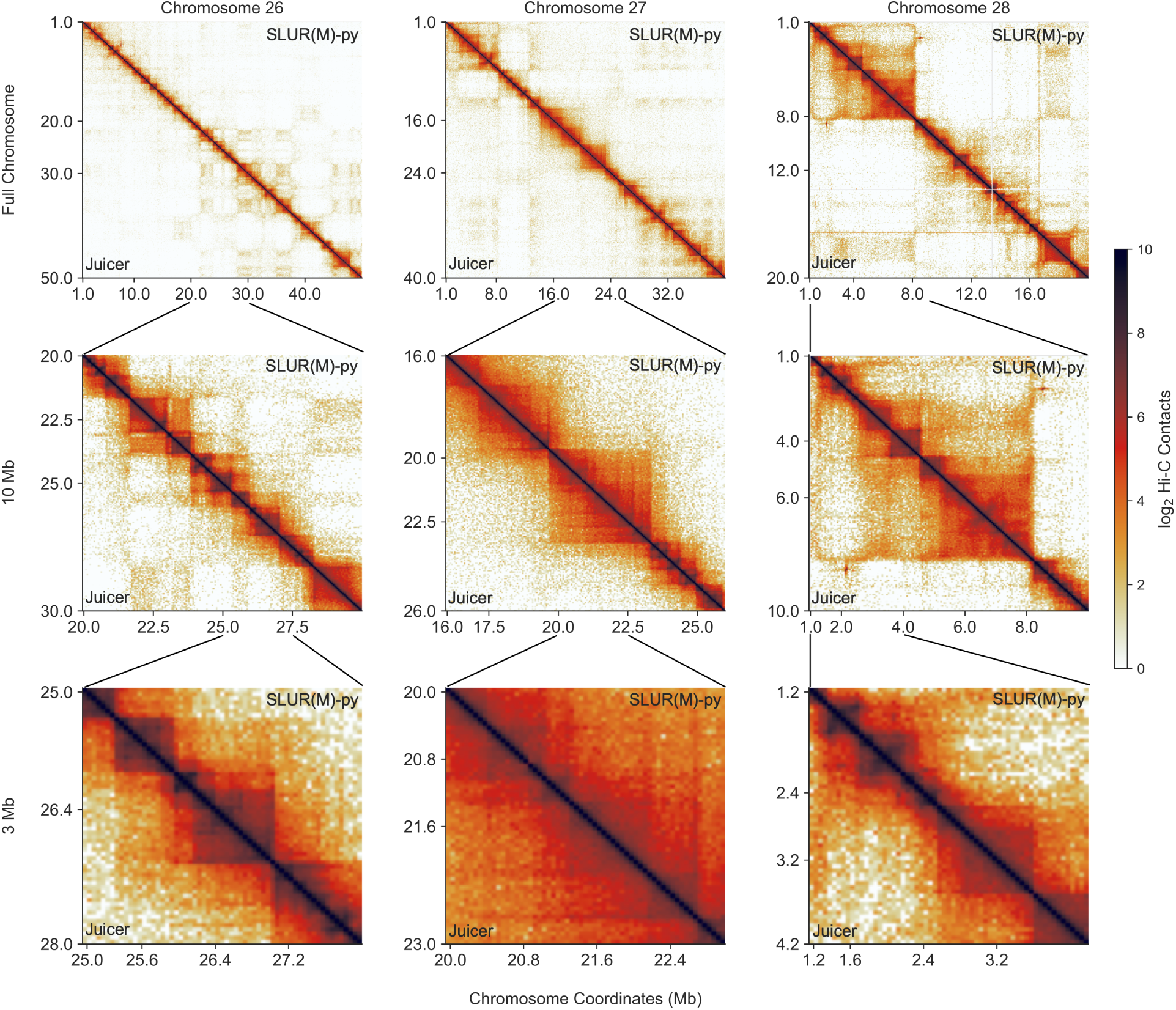
Comparison of Hi-C maps generated by Juicer and SLUR(M)-py pipelines. Example Hi-C maps for chromosomes 26, 27, and 28 (left to right, respectively) of the VERO genome, at scales of the full chromosome, 10 Mb, and 3 Mb (top to bottom, respectively). The Hi-C maps made from Juicer or SLUR(M)-py appear on the lower and upper diagonals, respectively. Orange to dark orange colors depict increasing contact frequency (log_2_) along the chromosomes.

**Figure S4:**
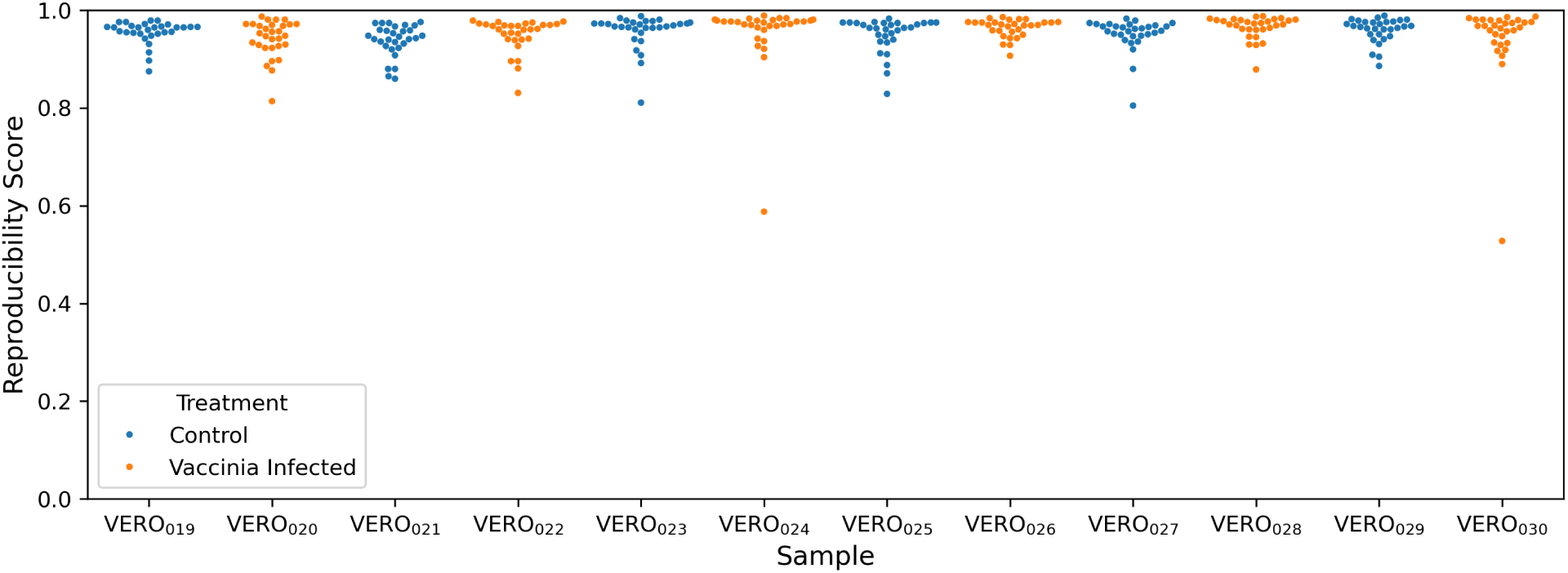
Reproducibility scores between Hi-C maps made by Juicer and SLUR(M)-py. Across Hi-C samples from Venu, et al. (2024), (x-axis) the spector reproducibility (y-axis) score was calculated between Hi-C maps (per chromosome) made by the Juicer pipeline and those processed by the SLUR(M)-py pipeline. Dots within swarm plots represent scores per chromosome for a given sample. Blue and orange colors denote control samples and those samples infected with vaccinia virus. Only two infected samples show a score below 0.80 for chromosome 19 (NC_023660.1).

**Figure S5:**
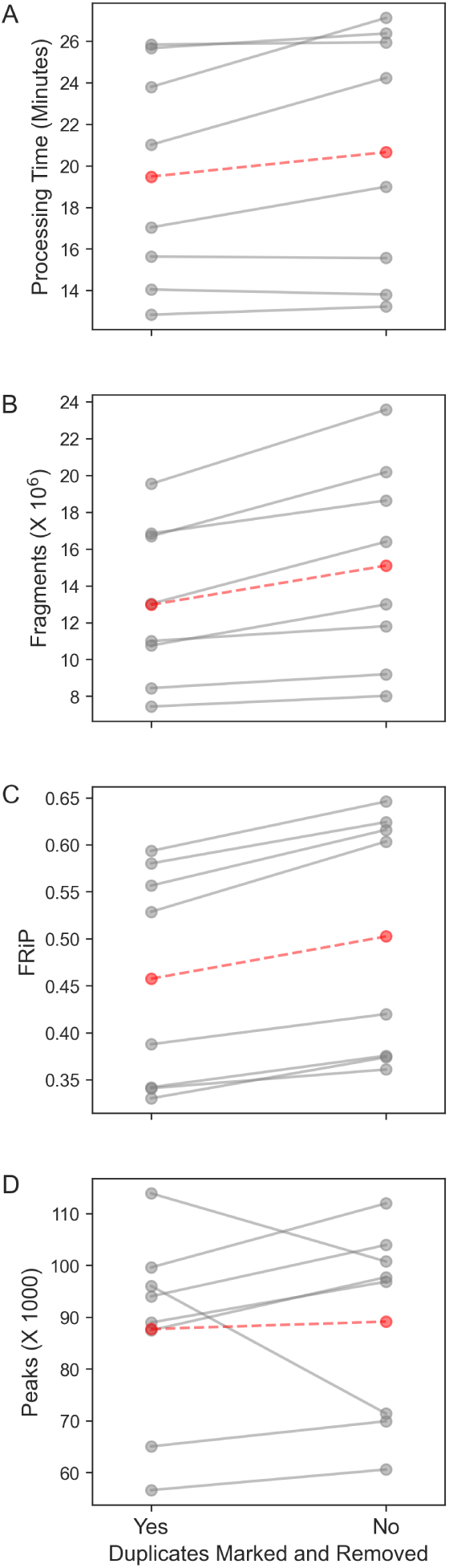
The effect of marking and removing duplicates on processing times, valid fragment counts, FRiP scores, and total peak counts across ATAC-seq experiments. **A**) The number of minutes (y-axis) for marking and removing duplicates, filtering alignments, and calling MACS3 across ATAC-seq experiments in the A549 cell line. **B**) The total number of valid fragments (y-axis) called from analysis. **C**) The fraction of reads in peaks (FRiP, y-axis) calculated by SLUR(M)-py. **D**) The number of detected peaks (via MACS3) in each ATAC-seq experiment (y-axis). The x-axis is shared in each panel and denotes the change in processing times, fragment counts, FRiP scores, and number of detected peaks (**A**, **B**, **C**, and **D** respectively) when including or excluding duplicate marking and removal. Red horizontal lines and dots mark the averages of each subplot.

**Figure S6:**
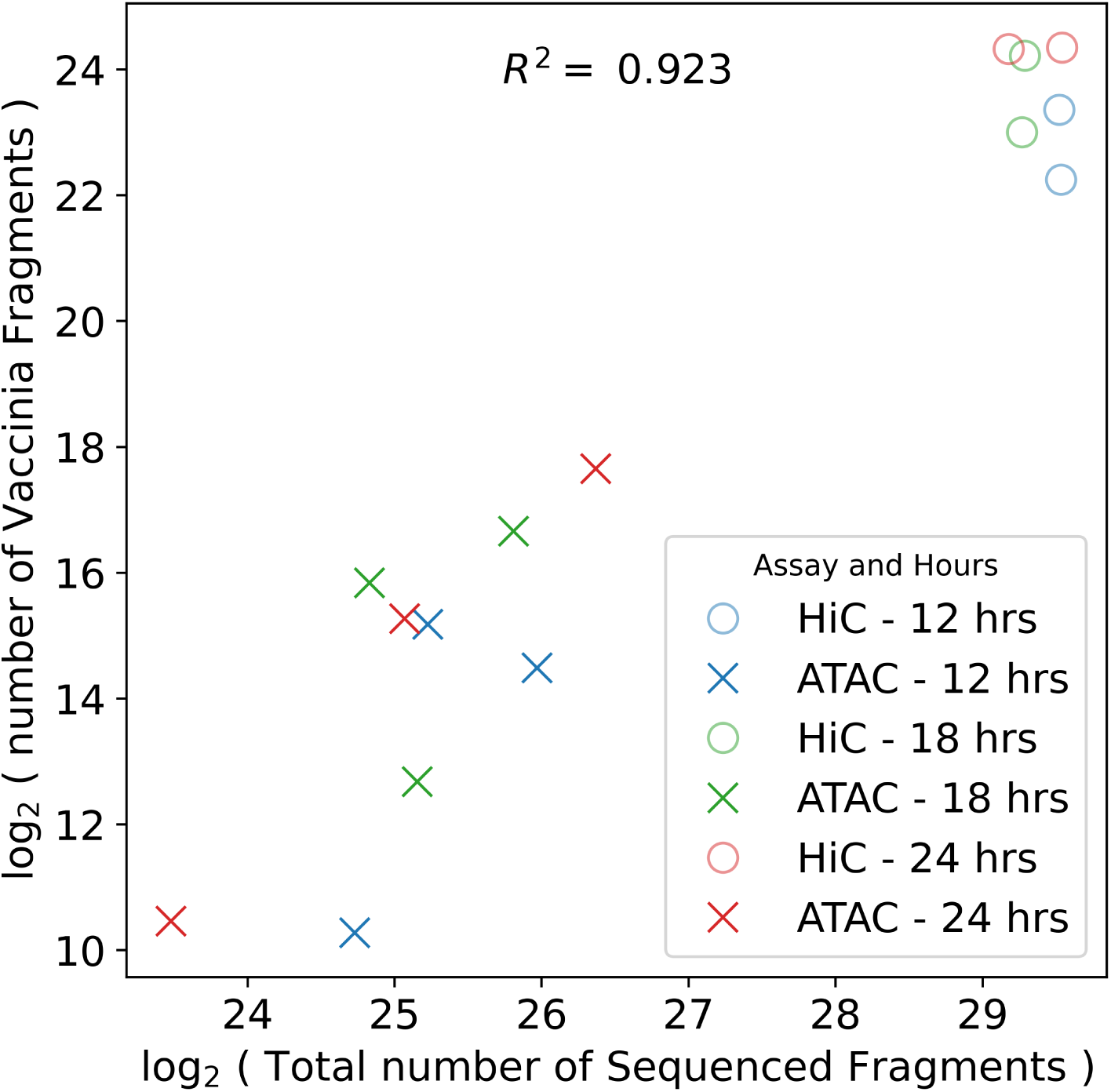
Counts of sequenced vaccinia fragments across ATAC-seq and Hi-C experiments. The count of sequenced fragments (y-axis) mapped to the vaccinia contig as a function of the total number of sequenced fragments (x-axis). The Xs and Os mark counts from an ATAC-seq or Hi-C experiment, respectively. Blue, green, and red colors denote the number of hours post infection. There is a significant linear relationship between the total identified vaccinia fragments and the number of sequenced reads in sequencing experiments on infected cells (*p*-value *<* 0.01, *R*^2^ = 0.923).

**Figure S7:**
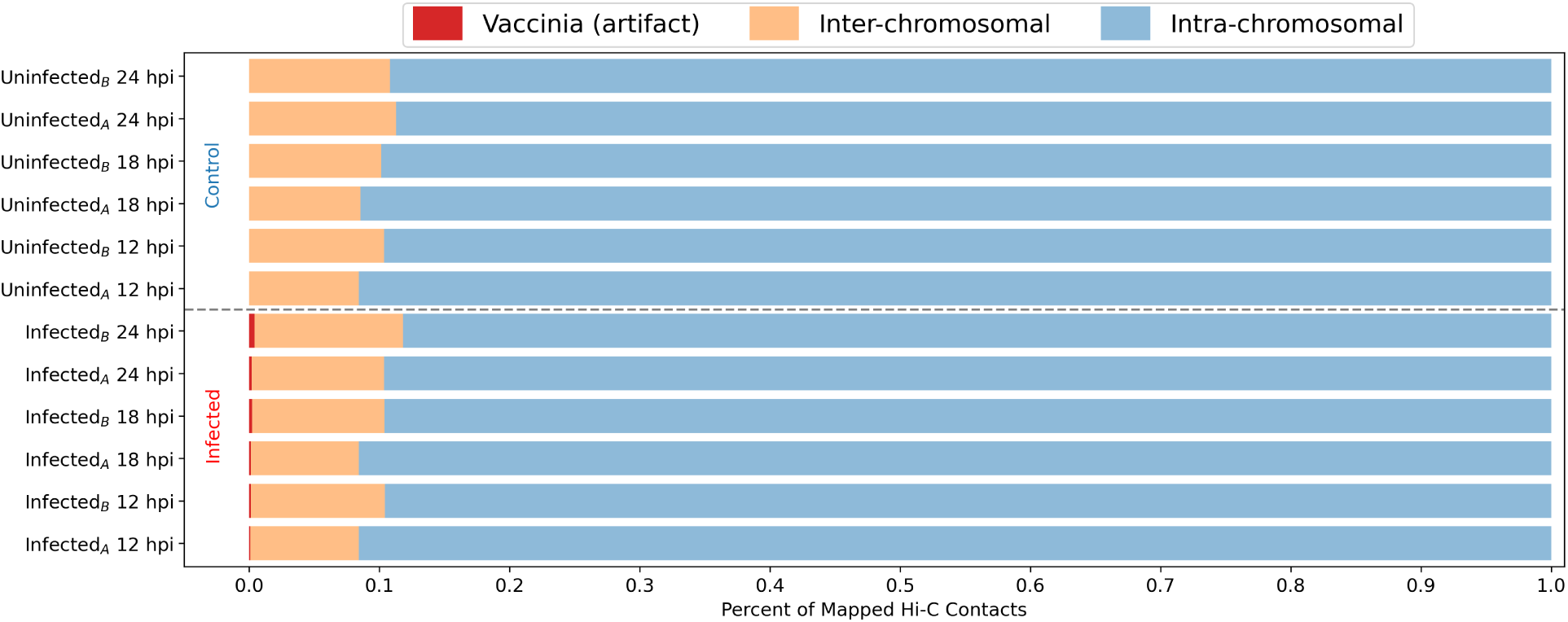
Inter- and intra-chromosomal contacts in control and infected Hi-C experiments. The portions (x-axis) of mapped Hi-C contacts are colored blue or orange across experiments (y-axis) for groups of paired-end reads mapping within or between chromosomes (respectively). Within infected experiments (bottom six experiments) the small portion of paired-end reads mapping between the genome and the vaccinia contig are marked in red. These events represent miss-ligation events between the genome and the viral DNA occurring post cross-linking within the cellular nucleus.

**Figure S8:**
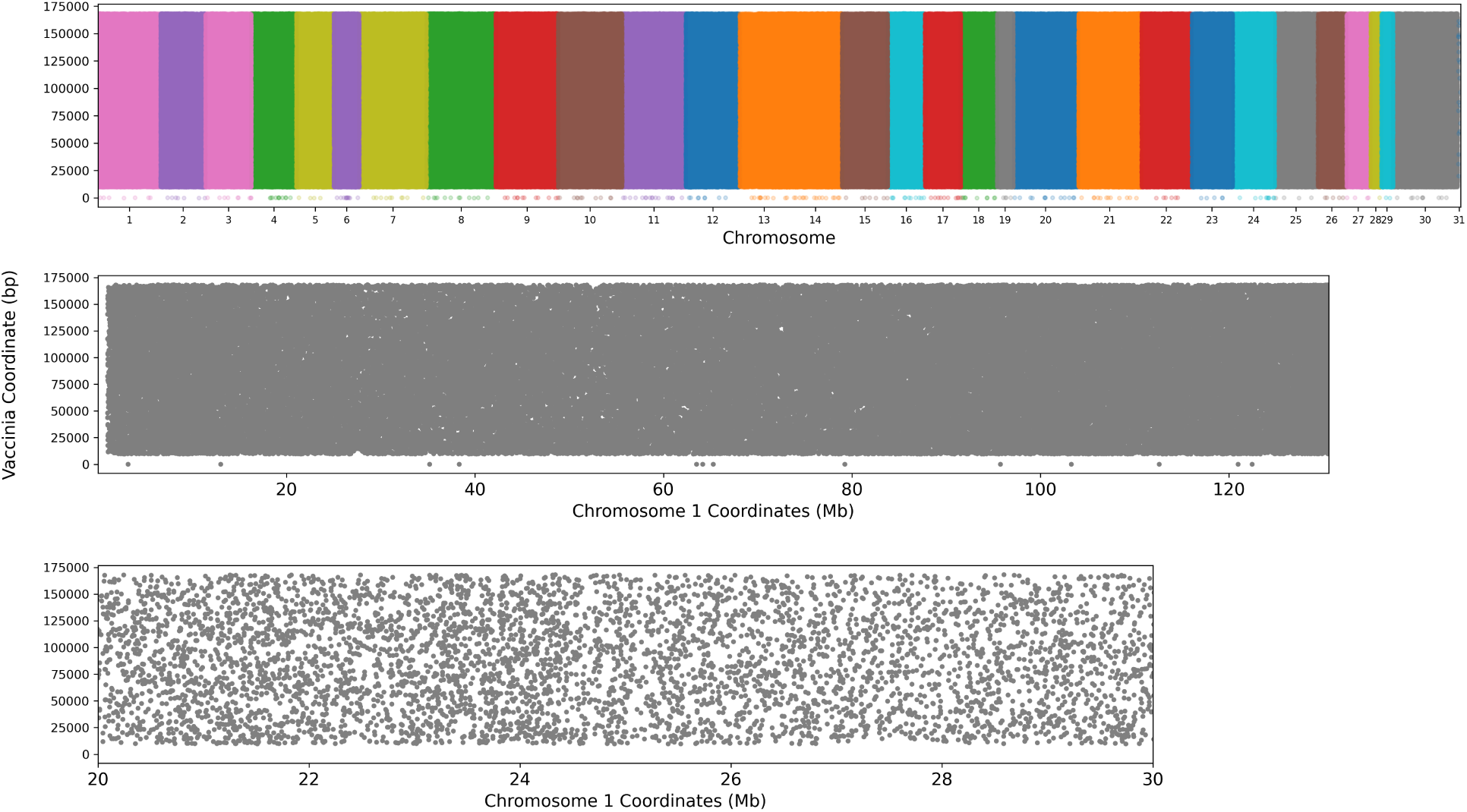
Mapping positions of artificial, miss-ligation events between vaccinia virus and genome. **Top**: Colored dots mark the locations of a ligation event between the double stranded vaccinia virus and the genome. The locations of these events along the vaccinia contig and the green monkey chromosome are marked along the y- and x-axis, respectively. **Middle**: The locations of artificial ligation events between the vaccinia virus and chromosome 1 of the Green Monkey genome. Each gray dot represents the positioning of a single ligation event between the genome (x-axis) and the viral contig (y-axis). **Bottom**: A 10 Mb region along chromosome 1 displaying ligation events and their locations between the genome (x-axis) and the viral contig (y-axis). At all three resolutions, genomic (**top**), chromosomal (**middle**), and 10 Mb (**bottom**), the ligation between genomic and viral DNA is uniformly distributed.

**Figure S9:**
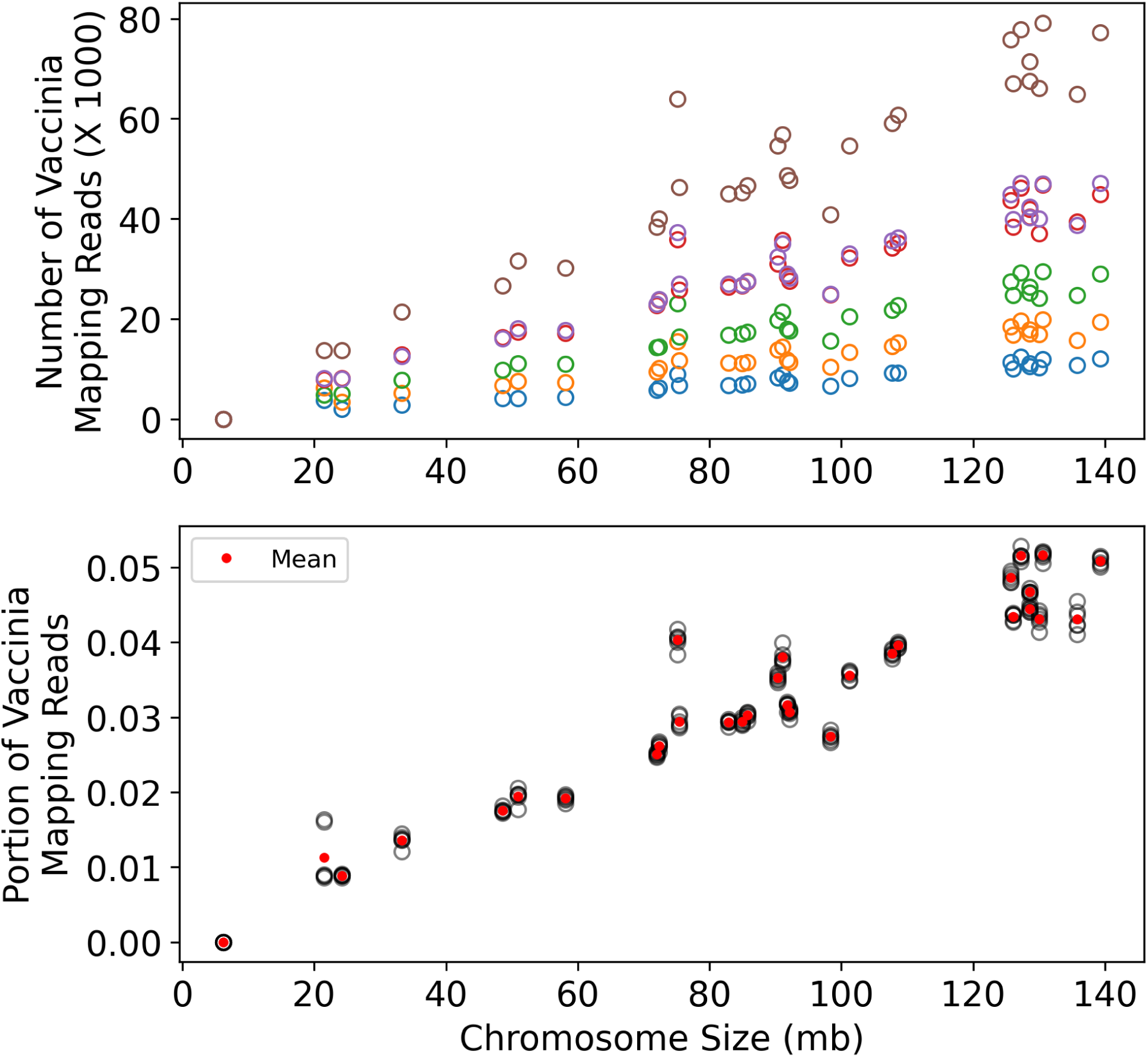
Counts of vaccinia mapped reads across the Green Monkey chromosomes. Top: Across replicated sequencing experiments (separated by colors) The number of identified read pairs mapping between the vaccinia contig and a given Green Monkey chromosome (y-axis) is plotted as a function of chromosome size. Bottom: The portions (y-axis, black circles) of vaccinia mapping reads as a function of chromosome size (x-axis). The mean portion across experiments is displayed via red dots.

**Figure S10:**
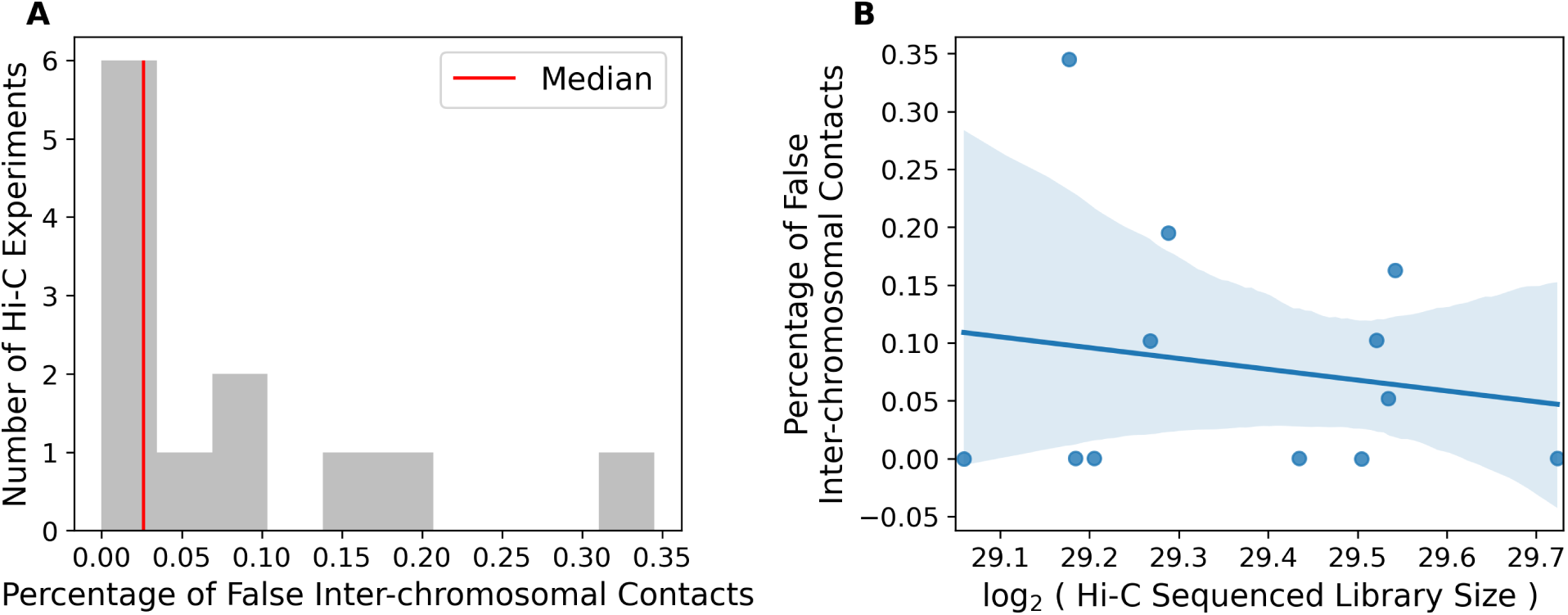
Percent of false positive inter-chromosomal contacts. **A**: Distribution of the percent of ligation events between the vaccinia contig and Green Monkey genome determined to be false positive inter-chromosomal contacts. The median percentage across experiments was 0.0261 (red vertical line). **B**: The percentage of false inter-chromosomal contacts (y-axis) versus the sequenced reads within each Hi-C library (x-axis). Blue line and shaded region represent a regression model and its 95% confidence interval relating the false inter-chromosomal contacts to the log_2_ of the read size. There is no significant linear relationship between the false positive percentage and number of reads (*p*-value *>* 0.05, *R*^2^ = 0.029).

**Figure S11:**
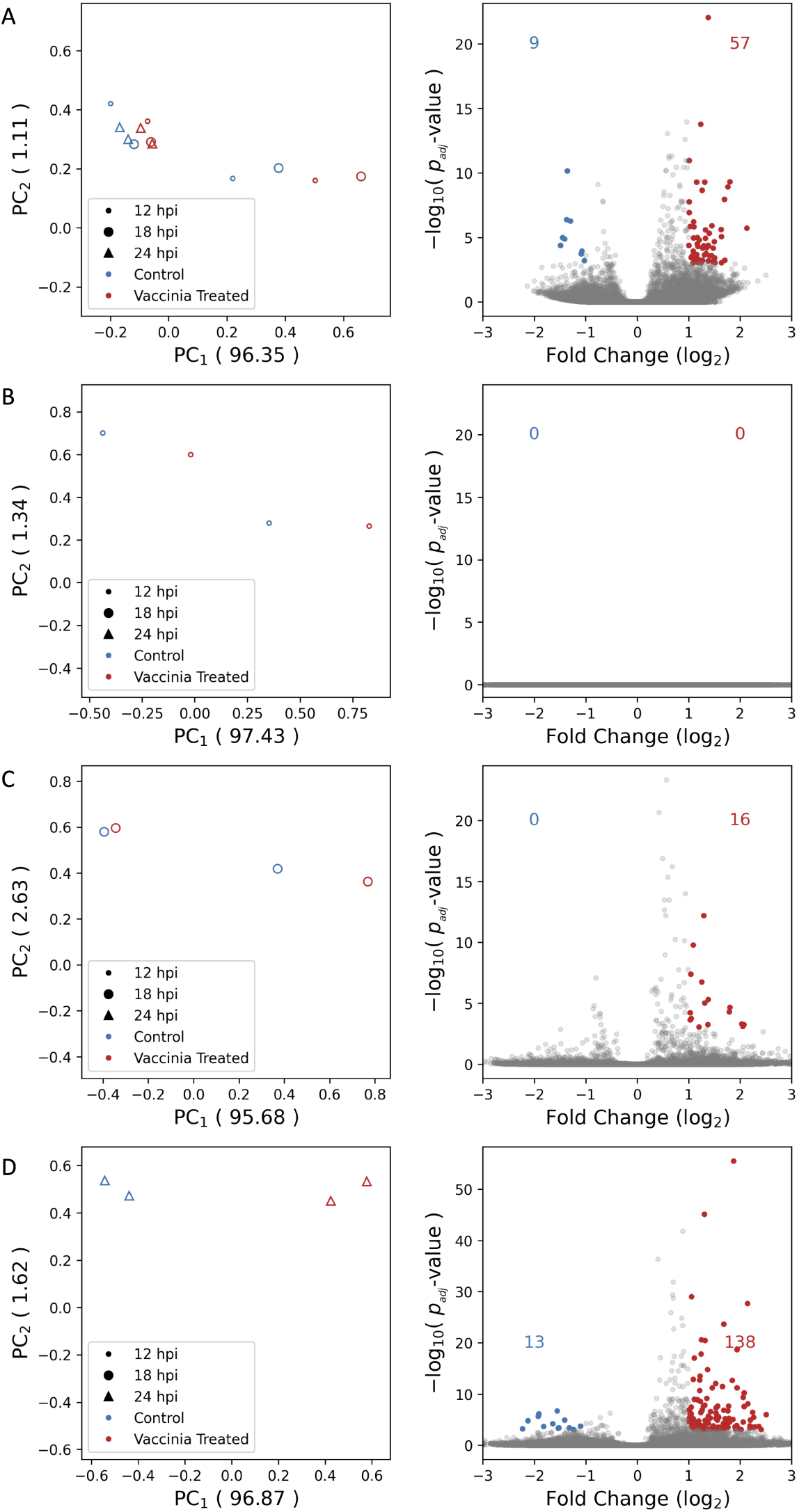
Analysis of inter-chromosomal contacts in VERO cells across vaccinia infection time-course. **A**) **Left**: the principal component analysis of 500 kb binned counts of contacts between chromosomes across vaccinia infected samples (red) and controls (blue) throughout time at 12, 18, and 24 hours post infection (hpi) (dots, circles, and triangles, respectively). **Right**: Differential analysis of inter-chromosomal contacts. Red and blue colors depict binned contacts between chromosomes increasing or decreasing due to vaccinia infection, respectively, across time. **B-D**) Principal component analysis (left) and differential analysis (right) of inter-chromosomal contacts at 12, 18, and 24 hpi.

**Table S1:** Processing, Mapping, and Peak Statistics of ATAC-seq Samples.

| Cell Line | Run Time | Paired-end Reads | Mapped (q > 30) | Peaks | Summits | FRiP | Genome Bases Covered ( percent ) | Source |
| --- | --- | --- | --- | --- | --- | --- | --- | --- |
| A549 | 0:19:17 | 16351381 | 11619607 | 89580 | 114555 | 0.5865 | 43693765 ( 1.402 ) | [40] |
| A549 | 0:22:28 | 21204135 | 10945957 | 90265 | 104479 | 0.3592 | 46231355 ( 1.483 ) | [40] |
| A549 | 0:17:48 | 16021189 | 8514473 | 64258 | 76240 | 0.3625 | 27657701 ( 0.887 ) | [40] |
| A549 | 0:44:35 | 33406352 | 16987737 | 92792 | 108354 | 0.3681 | 51768427 ( 1.661 ) | [40] |
| A549 | 0:22:23 | 17318402 | 7460950 | 55477 | 64813 | 0.4074 | 31944717 ( 1.025 ) | [40] |
| A549 | 0:36:13 | 21740996 | 20690406 | 99288 | 125802 | 0.5948 | 66678633 ( 2.139 ) | [40] |
| A549 | 0:27:50 | 21074428 | 13940998 | 113389 | 139868 | 0.6253 | 59522589 ( 1.909 ) | [40] |
| A549 | 0:35:11 | 24469297 | 17429501 | 86591 | 109579 | 0.621 | 65863808 ( 2.113 ) | [40] |
| VERO | 0:27:43 | 11125133 | 15688912 | 112610 | 144830 | 0.4859 | 79265606 ( 2.889 ) | [41] |
| VERO | 0:24:14 | 13919640 | 7743792 | 81198 | 102970 | 0.439 | 49251432 ( 1.795 ) | [41] |
| VERO | 0:55:23 | 24173485 | 30454312 | 132871 | 208144 | 0.4061 | 114774486 ( 4.183 ) | [41] |
| VERO | 0:34:59 | 18691465 | 12217888 | 100913 | 134291 | 0.308 | 69594515 ( 2.542 ) | [41] |
| VERO | 0:25:15 | 9492136 | 12803176 | 114561 | 147878 | 0.3904 | 95182764 ( 3.476 ) | [41] |
| VERO | 0:12:10 | 5848626 | 3948012 | 67851 | 71947 | 0.2171 | 29596890 ( 1.079 ) | [41] |
| VERO | 0:38:51 | 21299383 | 19135402 | 113191 | 152489 | 0.6799 | 60565293 ( 2.207 ) | [41] |
| VERO | 0:30:53 | 19644978 | 5754557 | 81310 | 98461 | 0.4512 | 37448317 ( 1.365 ) | [41] |
| VERO | 0:16:43 | 8547245 | 7805617 | 104767 | 127810 | 0.7415 | 47891003 ( 1.745 ) | [41] |
| VERO | 0:22:16 | 14905800 | 3326216 | 49718 | 60790 | 0.4342 | 22312359 ( 0.813 ) | [41] |
| VERO | 0:20:12 | 10941701 | 9714255 | 99634 | 126874 | 0.7038 | 47536681 ( 1.732 ) | [41] |
| VERO | 0:26:49 | 17601470 | 4288293 | 62048 | 73774 | 0.3361 | 29699129 ( 1.082 ) | [41] |
| VERO | 1:25:40 | 39831755 | 39798249 | 245706 | 354836 | 0.3814 | 110789009 ( 4.037 ) | [41] |
| VERO | 0:51:44 | 32864186 | 13174253 | 115644 | 156562 | 0.5307 | 51187658 ( 1.865 ) | [41] |
| VERO | 1:00:24 | 32288959 | 26392861 | 122195 | 166423 | 0.6018 | 61006806 ( 2.223 ) | [41] |
| VERO | 0:45:12 | 29424885 | 8528669 | 67641 | 88676 | 0.3599 | 28950015 ( 1.055 ) | [41] |
| VERO | 1:11:34 | 39334143 | 30132783 | 146057 | 201135 | 0.5834 | 71671681 ( 2.612 ) | [41] |
| VERO | 1:09:48 | 43316533 | 17979081 | 123494 | 169877 | 0.4162 | 55772205 ( 2.032 ) | [41] |

**Table S2:** Cell lines, Read Counts, and SLUR(M)-py Run Times of Hi-C data.

| Sample Name | Paired-end Reads | Run Time (HH:MM:SS) | Valid Hi-C Contacts | Fragmentation Enzyme | Cell Line | Source |
| --- | --- | --- | --- | --- | --- | --- |
| A549<sub>000</sub> | 130305900 | 01:03:45 | 78143020 | HindIII | A549 | ENCLB222WYT |
| A549<sub>001</sub> | 121585833 | 00:58:13 | 74012587 | HindIII | A549 | ENCLB571HTP |
| A549<sub>002</sub> | 254857397 | 04:48:50 | 172165621 | MboI | A549 | ENCLB571GEP |
| A549<sub>021</sub> | 727353489 | 10:39:44 | 322065131 | Arima | A549 | This study |
| HepG2<sub>003</sub> | 815725922 | 08:51:10 | 739526295 | HindIII | HepG2 | ENCLB022KPF |
| HepG2<sub>004</sub> | 885833793 | 10:01:18 | 771404739 | HindIII | HepG2 | ENCLB625TGE |
| IMR90<sub>005</sub> | 252725458 | 01:59:11 | 173539638 | MboI | IMR90 | ENCLB476ZIC |
| IMR90<sub>006</sub> | 276809208 | 02:20:03 | 190530643 | MboI | IMR90 | ENCLB512HIP |
| K562<sub>007</sub> | 591854553 | 06:48:44 | 363306188 | MboI | K562 | ENCLB470UUQ |
| K562<sub>008</sub> | 456757799 | 03:08:49 | 305186793 | MboI | K562 | ENCLB941YRT |
| VERO<sub>019</sub> | 559483569 | 12:29:35 | 265274243 | Arima | VERO | [41] |
| VERO<sub>020</sub> | 777613496 | 10:12:05 | 274526191 | Arima | VERO | [41] |
| VERO<sub>021</sub> | 761567093 | 09:46:21 | 344542129 | Arima | VERO | [41] |
| VERO<sub>022</sub> | 646594007 | 08:17:01 | 280411179 | Arima | VERO | [41] |
| VERO<sub>023</sub> | 886507172 | 12:40:27 | 544176258 | Arima | VERO | [41] |
| VERO<sub>024</sub> | 770418243 | 11:01:13 | 459796419 | Arima | VERO | [41] |
| VERO<sub>025</sub> | 725697285 | 10:50:32 | 453580308 | Arima | VERO | [41] |
| VERO<sub>026</sub> | 781644457 | 11:01:34 | 446320262 | Arima | VERO | [41] |
| VERO<sub>027</sub> | 610190915 | 08:43:18 | 384589159 | Arima | VERO | [41] |
| VERO<sub>028</sub> | 655739644 | 09:24:57 | 392837985 | Arima | VERO | [41] |
| VERO<sub>029</sub> | 619039813 | 08:52:24 | 410112037 | Arima | VERO | [41] |
| VERO<sub>030</sub> | 607026371 | 08:23:21 | 365787228 | Arima | VERO | [41] |

